# Inhibition of early-acting autophagy genes in *C. elegans* neurons improves protein homeostasis, promotes exopher production, and extends lifespan via the ATG-16.2 WD40 domain

**DOI:** 10.1101/2022.12.12.520171

**Authors:** Yongzhi Yang, Meghan Lee Arnold, Elizabeth H. Choy, Caitlin M. Lange, Karie Poon, Michael Broussalian, Ling-Hsuan Sun, Tatiana M. Moreno, Anupama Singh, Monica Driscoll, Caroline Kumsta, Malene Hansen

## Abstract

While autophagy is key to maintain cellular homeostasis, tissue-specific roles of individual autophagy genes are less understood. To study neuronal autophagy *in vivo*, we inhibited autophagy genes specifically in *C. elegans* neurons, and unexpectedly found that knockdown of early-acting autophagy genes, i.e., involved in formation of the autophagosome, except for *atg-16.2*, decreased PolyQ aggregates and increased lifespan, albeit independently of the degradation of autophagosomal cargo. Neuronal aggregates can be secreted from neurons via vesicles called exophers, and we found that neuronal inhibition of early-acting autophagy genes *atg-7* and *lgg-1/Atg8*, but not *atg-16.2* increased exopher formation. Moreover, *atg-16.2* mutants were unable to form exophers, and *atg-16.2* was required for the effects of early autophagy gene reduction on neuronal PolyQ aggregation, exopher formation, and lifespan. Notably, neuronal expression of full-length ATG-16.2 but not ATG-16.2 without a functional WD40 domain, important for non-canonical functions of ATG16L1 in mammalian cells, restored these phenotypes. Collectively, we discovered a specific role for *C. elegans* ATG-16.2 and its WD40 domain in exopher biogenesis, neuronal proteostasis, and lifespan determination, highlighting a possible role for non-canonical autophagy functions in both exopher formation and in aging.

## INTRODUCTION

Macroautophagy (hereafter referred to as autophagy) is an intracellular recycling process by which cytosolic material or cargo is subjected to lysosomal degradation, referred to as canonical autophagy. Autophagy plays important roles in numerous late-onset diseases including neurodegenerative disorders and has been directly linked to aging in multiple model organisms including the nematode *C. elegans*^1^. In this organism, RNA interference (RNAi) of multiple autophagy genes during adulthood abrogates lifespan extension in long-lived mutants, suggesting autophagy is required for longevity. In contrast, RNAi inhibition of autophagy genes in wild-type (WT) animals typically has small or no effects on lifespan^1^, indicating that basal autophagy is not limiting for normal aging. Still, the tissue-specific contributions of autophagy to organismal fitness and lifespan extension remain unclear. In particular, the role of autophagy genes in neurons is of special interest since neuronal signaling plays a key role in several longevity paradigms^2^, and remains to be thoroughly investigated in *C. elegans,* where neurons are generally refractory to RNAi interference^3-6^.

Autophagy is a multi-step process that includes at least five steps: (i) initiation, (ii) nucleation and formation of a double-membrane structure (phagophore), (iii) phagophore elongation and enclosure to form an autophagosome that incorporates cargo such as misfolded and aggregated proteins, (iv) autolysosome formation by fusion of the autophagosome and lysosome, and finally (v) cargo degradation in the autolysosome^7^. Multiple conserved autophagy-related (Atg) genes function in canonical autophagy, and genes acting in steps before autophagosome and lysosome fusion are considered early-acting autophagy genes, whereas genes functioning in autolysosome formation and cargo degradation can be considered late-acting autophagy genes^8^. During canonical autophagy, activation of the ATG1/ULK1 complex initiates the formation of the phagophore, which is further facilitated by the phosphatidylinositol (PtdIns) 3-kinase (PtdIns3k) complex and PtdIns 3-phosphate (PtdIns3P)-binding complexes. Phagophore elongation requires two conjugation complexes, the ATG5-ATG12/ATG16 complex and the ATG8 complex. These two complexes promote the conjugation of ATG8 to phosphatidylethanolamine (PE) at both the inner and outer membrane of the growing double-membrane phagophore^9^. ATG8 is the most common reporter for pre-autophagosomes and closed autophagosomes^10,11^.

In recent years, emerging studies in mammalian tissue cultures have shown that in addition to their roles in canonical autophagy, some early-acting autophagy genes also have functions in other cellular processes, such as phagocytosis and secretion. In these non-canonical activities, ATG8 is conjugated to single-membrane vesicles by the ATG5-ATG12/ATG16 complex^12-14^. While it remains underexplored how the ATG5-ATG12/ATG16 complex differentiates between double- and single-membraned vesicles, the WD40 domain in the C terminus of ATG16L1 is specifically required for its non-canonical function in LC3-associated phagocytosis (LAP)^13^, highlighting this domain of ATG16L1 as a key molecular entity that can separate its canonical and non-canonical functions, at least in phagocytosis. While one study has indicated a role for the WD40 domain of ATG16L1 in preventing Alzheimer’s disease in mice^15^, non-canonical functions for autophagy genes have primarily been studied in mammalian cell culture, and its relevance has yet to be studied in the context of organismal aging.

To better understand the neuronal role of autophagy genes in neuronal proteostasis, we used RNAi to inhibit autophagy genes in individual tissues of adult *C. elegans.* Following neuronal inhibition of early-acting, but not late-acting autophagy genes, we observed a surprising decrease in neuronal aggregates of proteins with expanded polyglutamine (PolyQ) stretches, a model for Huntington’s disease, and an increase in the secretion of vesicles, called exophers, which was accompanied with an extended lifespan. Importantly, these phenotypes were *atg-16.2* dependent and could be rescued by pan-neuronal expression of ATG-16.2, an ortholog of mammalian ATG16L1, provided it included an intact WD40 domain. Our studies identified a differential role for early and late-acting autophagy genes in *C. elegans* neurons and discovered that ATG-16.2 and its highly conserved WD40 domain are required for exopher biogenesis, for maintaining neuronal proteostasis, and for lifespan extension, highlighting that, in this model, non-canonical agy gene functions may be critical for tissue- and organismal fitness via alternative ory pathways for disposal of misfolded proteins.

## RESULTS

### Neuronal inhibition of early-acting autophagy genes improves neuronal proteostasis and extends lifespan in *C. elegans*

*C. elegans* neurons are generally refractory to RNAi because they lack the RNA channel/transporter protein SID-1 required for RNA uptake^3-6^. Previous studies investigating autophagy genes by RNAi approaches have therefore not fully assessed the role of autophagy genes in neurons. To address this limitation, we constructed a new *C. elegans* strain capable of RNAi in neurons only by expressing SID-1 under the pan-neuronal *rgef-1* promoter in *sid-1(-)* loss-of-function mutants, similar to previous studies16. We validated this strain in multiple ways, including by showing that the *rgef-1* promoter remained expressed in neurons into late adulthood (**Figure S1a**), and that RNAi effects were specific and selective to neurons in this strain (**Figure S1b-d**), which displayed a normal lifespan (in 14 out of 16 experiments; P < 0.01, log-rank, **Table S1**).

Since whole-body reduction (i.e., primarily in non-neuronal tissues) of all tested autophagy genes increased the aggregation of YFP-tagged Polyglutamine (PolyQ) aggregates in *C. elegans* neurons (**Figure S2a**)^17-19^, we asked if neuronal aggregates was similarly affected when autophagy genes were reduced in neurons only [all of the analyzed genes are expressed in *C. elegans* neurons^20^]. While no changes were observed in PolyQ aggregation in *sid-1* mutants or upon neuronal expression of SID-1 (**Figure S2b**), we unexpectedly observed that neuronal-only knockdown of autophagy genes with functions in early-versus late-acting steps of the process (**Figure 1A**) showed noticeable differences in neuronal PolyQ aggregation. Specifically, neuronal-only inhibition of genes with functions early in the pathway (i.e., *unc-51/Atg1, atg-13, bec-1/Beclin1, atg-9*, *atg-7, atg-4.1,* and *lgg-1/Atg8)*, decreased neuronal PolyQ aggregate load, whereas knockdown of late-acting autophagy genes (i.e., *cup-5, epg-5, vha-13, vha-15*, and *vha-16*) had no effect or increased the number of neuronal PolyQ aggregates (**Figure 1B-C**), suggesting a differential effect of early versus late autophagy gene knockdown on neuronal proteostasis.

**Figure 1.**
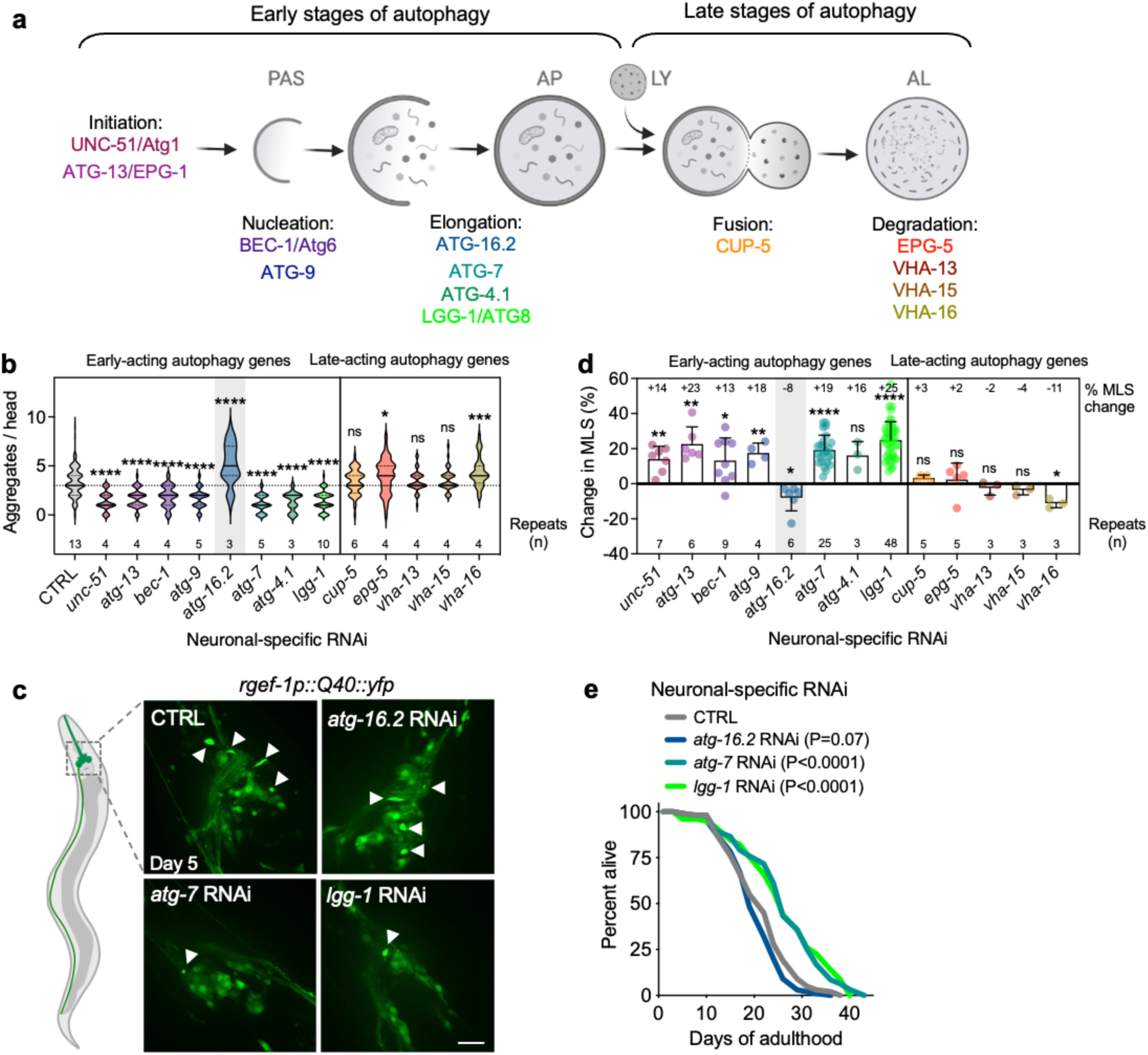
With exception of *atg-16.2*, neuronal inhibition of early-acting autophagy genes reduces neuronal PolyQ aggregation and extends lifespan. **(a)** Schematic diagram showing early- and late-acting genes in the macroautophagy (here referred to as autophagy) process investigated in this study (see introduction for additional information). (**b**) Number of neuronal PolyQ aggregates in day 5 animals capable of neuronal-only RNAi (*sid-1; rgef-1p::sid-1; rgef-1::Q40::yfp)* and fed bacteria expressing empty vector (grey) or dsRNA of the listed autophagy-related genes from hatching. Violin plots of data from indicated number of repeats (n), each with N > 7 animals with solid line indicating median and dashed lines indicating quartiles. ns P > 0.05, *P < 0.05, ***P < 0.001, ****P < 0.0001, by one-way ANOVA. (**c**) Representative images of nerve-ring neurons of day 5 animals capable of neuronal-only RNAi (*sid-1; rgef-1p::sid-1; rgef-1::Q40::yfp)* and fed bacteria expressing empty vector (CTRL), *atg-16.2, atg-7,* or *lgg-1/Atg8* dsRNA from hatching on with arrowheads indicating PolyQ aggregates. Scale bar: 20 µm. (**d**) Average mean lifespan change (% MLS change, indicated) in animals capable of neuronal-only RNAi (*sid-1; rgef-1p::sid-1)* fed bacteria expressing dsRNA of the listed autophagy-related genes compared to control. All lifespan data were pooled irrespective of RNAi initiation either from hatching on or starting in adult animals in the indicated 3-9 experimental repeats. Error bars indicate s.d. ns P > 0.05, *P < 0.05, **P < 0.01, ***P < 0.001, ****P < 0.0001, by one-sample t-test compared to hypothetical mean of 0. See **Table S2** for details of individual lifespan experiments. (**e**) Lifespan analyses of animals capable of neuronal-only RNAi (*sid-1; rgef-1p::sid-1 + rgef-1p::gfp*) fed bacteria expressing empty vector (CTRL), *atg-16.2*, *atg-7,* or *lgg-1/Atg8* dsRNA from hatching on. Statistical significance determined by log-rank test. See **Table S2** for details and repeats.

Since neuronal gene function can affect aging^2^, we also tested the effect of neuronal-only knockdown of autophagy genes on lifespan. Strikingly, we observed a similar, differential pattern for lifespan, i.e., extended lifespan was observed for whole-life or adult-only, neuronal-only inhibition of early-acting genes, including *atg-7* (in 20 out of 22 experiments; P < 0.01, log-rank,) and *lgg-1*/Atg8 (in 41 out of 46 experiments; P < 0.01, log-rank,), whereas reduction of genes with functions in later stages were incapable of doing so, and *vha-16* inhibition even shortened lifespan (**Figure 1d** **and Table S2**). Moreover, whole-life, neuronal-only RNAi against *atg-7* or *lgg-1/Atg8* also extended lifespan in short-lived animals expressing neuronal PolyQ aggregates (**Figure S2c, Table S3**), indicating that the decreased aggregation load conferred by neuronal inhibition of autophagy genes correlates with consequential effects on longevity. Interestingly, neuronal-only RNAi knockdown of early-acting gene *atg-16.2* behaved differently; we observed significant increase in PolyQ aggregates (**Figure 1b-c**), and no lifespan extension in any of 6 experiments (**Figure 1d-e** **and Table S2**), making it an exception to other early-acting autophagy genes, which all showed beneficial effects following neuronal knockdown.

We next used *atg-7* or *lgg-1/Atg8* RNAi clones as representative early-acting autophagy genes and tested the effects of neuronal-only knockdown on healthspan measures, and found that these animals displayed unchanged pharyngeal pumping, swimming ability, and progeny production (**Figure S3a-c**). Since defects in ciliated sensory neurons increase lifespan in *C. elegans*^21^, we also tested if neuronal-only RNAi against *atg-7* or *lgg-1/Atg8* affected the morphology of ciliated sensory neurons in dye-filling assays, to document that, day 5 animals subjected to RNAi displayed normal dye-filling phenotypes compared to control animals (**Figure S3d**). Our data suggest that defects in ciliated sensory neurons are not responsible for lifespan extension after neuronal knockdown of *atg-7* or *lgg-1/Atg8*. To further test whether neuronal autophagy gene reduction affected neuronal physiology, we also counted the neuronal branches that grow out from mechanoreceptor neurons, such as ALML/R, PLML/R, AVM, and PVM, at older age; a phenotype associated with neuronal dysfunction^22,23^. While animals treated with empty vector control showed branches originating from ALM and PLM mechanoreceptor neurons at an old age (day 15), this increase was not observed in animals subjected to neuronal-only RNAi against *atg-7* or *lgg-1/Atg8* (**Figure S3e**), suggesting that inhibition of early autophagy genes improves age-related decline in neuronal morphology.

Taken together, we observed that neuronal knockdown of an extensive panel of early-acting autophagy genes, except for *atg-16.2*, cell-autonomously reduced neuronal PolyQ aggregation and extended lifespan. The benefits induced by neuronal-only inhibition of at least *atg-7* and *lgg-1/Atg8* came with no significant effects on muscle-related healthspan measures, no impairment of progeny production, but did improve neuronal morphology. These results demonstrate a differential role for early-vs. late-acting acting autophagy genes in *C. elegans* neurons, and this differential effect indicates that the observed improvement in neuronal proteostasis and lifespan extension associated with their knockdown may not be a result of impairing lysosomal degradation per se.

### Neuronal inhibition of both early- and late-acting autophagy genes impairs autophagy

Considering the differential effects of inhibiting early- and late-acting autophagy genes in neurons, we next tested the effects of inhibiting autophagy genes on markers of autophagy in our neuronal-only RNAi strains, which had unchanged autophagy dynamics (**Figure S4a-b**). First, we used animals stably expressing GFP::LGG-1/Atg8, a reporter commonly used for the assessment of pre-autophagosomal phagophores and autophagosomes^24^, from a pan-neuronal *rgef-1* promotor^25^. We subjected these animals to neuronal RNAi of the full panel of autophagy genes and found that RNAi clones for early-acting autophagy genes, but not late-acting autophagy genes, generally decreased GFP::LGG-1 punctae counts (**Figure 2a**), consistent with early-acting autophagy genes being important for the formation of autophagosomes^26^. As a control, we repeated this experiment in neuronal-only RNAi strains expressing GFP::LGG-1(G116A) in neurons^18^. GFP::LGG-1(G116A) is lipidation deficient and lacks Glycine 116, which is important for conjugation to autophagosome membranes; as a consequence, GFP::LGG-1(G116A) should be evenly distributed in the cytosol. However, a low level of GFP-positive punctae were observed in GFP::LGG-1(G116A)-expressing animals (**Figure 2b**)^18^. While the nature of these GFP::LGG-1(G116A) structures has not been fully characterized, they may represent protein aggregates27. Interestingly, GFP::LGG-1(G116A) expressing animals subjected to neuronal-only inhibition of all early-acting autophagy genes, along with two late-acting genes, displayed a significant increase in the numbers of GFP::LGG-1(G116A) punctae (**Figure 2b**), consistent with an accumulation of protein aggregates from an autophagy block. To directly assess autophagy status in neurons, we used neuronal RNAi strains expressing the autophagy receptor SQST-1::GFP, a substrate for degradative autophagy^17^. Neuronal inhibition of most early- and several late-acting genes caused SQST-1::GFP accumulation in the head (**Figure 2c**), indicating that these RNAi treatments generally resulted in a block of canonical, degradative autophagy of at least SQST-1 receptor-related substrates.

**Figure 2.**
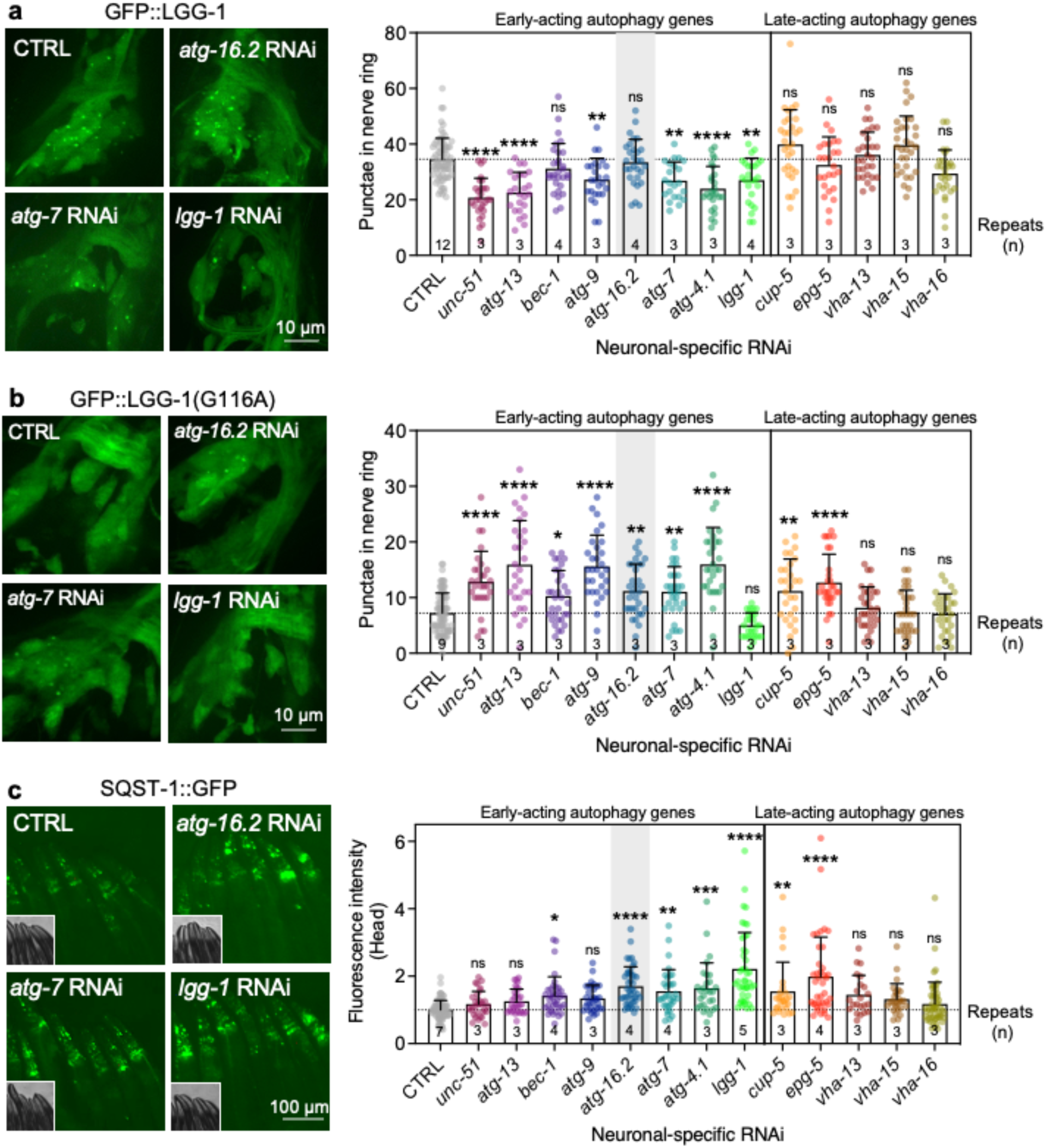
Neuronal inhibition of early-acting autophagy genes, including *atg-16.2*, impairs autophagy. (**a**) Representative images of nerve-ring neurons of animals capable of neuronal-only RNAi and expressing GFP::LGG-1/Atg8 pan-neuronally (*sid-1; rgef-1p::sid-1; rgef-1::gfp::lgg-1)* on day 1 of adulthood fed bacteria expressing empty vector (CTRL), *atg-16.2*, *atg-7,* or *lgg-1/Atg8* dsRNA from hatching on. Summary of average number of GFP::LGG-1/Atg8 punctae in nerve-ring neurons of these animals as well as in animals fed dsRNA of additional autophagy-related genes from hatching on. Data pooled from indicated number of experiments (n), with N=24-74 animals. Error bars indicate s.d. ns P > 0.05, *P < 0.05, **P < 0.01, ***P < 0.001, ****P < 0.0001, by one-way ANOVA. (**b**) Representative images of nerve-ring neurons of animals capable of neuronal-only RNAi and expressing non-lipidated GFP::LGG-1/Atg8(G116A) pan-neuronally (*sid-1; rgef-1p::sid-1; rgef-1::gfp::lgg-1(G116A))* on day 1 of adulthood fed bacteria expressing CTRL, *atg-16.2*, *atg-7,* or *lgg-1/Atg8* dsRNA from hatching on. Summary of average GFP::LGG-1(G116A) punctae in nerve-ring neurons as in **a**. Data pooled from indicated number of experiments (n), with N=26-91 animals. Error bars indicate s.d. ns P > 0.05, *P < 0.05, **P < 0.01, ***P < 0.001, ****P < 0.0001, by one-way ANOVA. (**c**) Representative images of head regions of animals capable of neuronal-only RNAi and expressing the autophagy receptor SQST-1/p62::GFP (*sid-1; rgef-1p::sid-1; sqst-1p::sqst-1::gfp)* on day 1 of adulthood fed bacteria expressing CTRL, *atg-16.2*, *atg-7,* or *lgg-1/Atg8* dsRNA from hatching on. Summary of GFP fluorescence intensity in head region as in **a-b**. Data pooled from indicated number of experiments (n), with N=22-90 animals. Error bars indicate s.d. ns P > 0.05, *P < 0.05, **P < 0.01, ***P < 0.001, ****P < 0.0001, by one-way ANOVA.

These results imply that early-autophagy gene inhibition can extend lifespan while autophagy activity or cargo degradation is impaired and indicate that the reduced PolyQ aggregation load and longevity phenotypes may be independent of canonical autophagy.

### Neuronal inhibition of *atg-7* or *lgg-1/Atg8,* but not *atg-16.2*, increases exopher biogenesis

Similar to human neurodegenerative diseases, neurons in *C. elegans* can rid themselves of toxic protein aggregates via biogenesis of large vesicles called exophers that jettison cytosolic material including PolyQ aggregates from neurons into surrounding tissues^28^. Since whole-body RNAi against early-acting autophagy genes *bec-1/Beclin1, atg-7*, and *lgg-1/lgg-2/Atg8*, increases exopher generation^28^, we tested the effect of neuronal knockdown of *atg-7* and *lgg-1/Atg8,* which decreased neuronal PolyQ aggregates (**Figure 1b-c**), on exopher biogenesis in ALMR neurons using an mCherry reporter expressed from the *mec-4* promoter (**Figure 3a**)^28,29^. While baseline exopher numbers were unchanged in neuronal RNAi strains (**Figure S4c**), we found that neuronal knockdown of *atg-7* and *lgg-1/Atg8* significantly increased exopher formation (**Figure 3b-c**). As expected, neuronal reduction of *atg-7* and *lgg-1/Atg8* also extended lifespan in the mCherry-reporter background (**Table S3).** In contrast, neuronal reduction of *atg-16.2*, which increased neuronal aggregation of PolyQ proteins (**Figure 1b-c**) and did not extend lifespan (**Figure 1d**), did not increase exopher generation (**Figure 3d**). Collectively, these results indicate correlation between reduced PolyQ aggregates, increased exopher formation, and lifespan extension observed in animals with neuronal inhibition of early-autophagy genes *atg-7* and *lgg-/*Atg8, unlike neuronal knockdown of *atg-16.2* which does not induce any of the aforementioned phenotypes.

**Figure 3.**
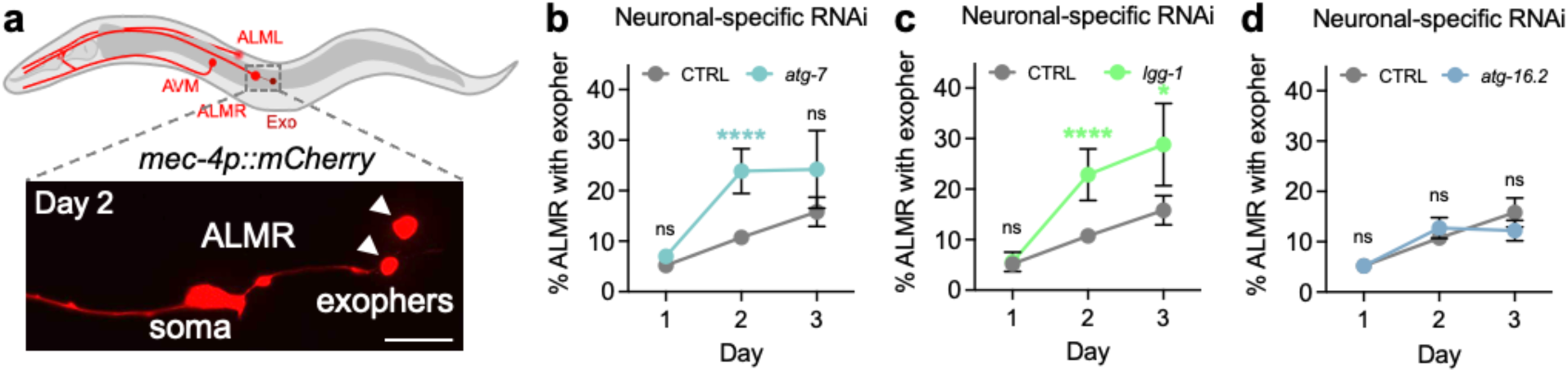
Neuronal inhibition of autophagy genes *atg-7* and *lgg-1/Atg8*, but not *atg-16.2,* increases exopher formation. **(a)** Exophers originating from the ALMR neuron in animals capable of neuronal-only RNAi and expressing *mec-4p*::mCherry (*sid-1; rgef-1p::sid-1; mec-4p::mCherry*). Representative picture showing two exophers (arrowheads) and the ALMR soma on day 2. Scale bar: 20 µm. (**b-d**) Quantification of percent of animals capable of neuronal-only RNAi (*sid-1; rgef-1p::sid-1; mec-4p::mCherry*) with exophers on day 1-3 of adulthood when fed bacteria expressing empty vector (CTRL), *atg-7* (**b**), *lgg-1/Atg8* (**c**), or *atg-16.2* (**d**) dsRNA from hatching. Error bars indicate mean and s.d. of n=5-8 experiments, each with N=26-61 animals. ns P > 0.05, *P < 0.05, ****P < 0.0001, by Cochran-Mandel Hansel test.

### *atg-4.1* and *atg-16.2* both impair autophagy, but neuronal inhibition of *atg-7* or *lgg-1/Atg8* does not lead to benefits in *atg-16.2* mutants

Considering that *atg-16.2* inhibition in neurons produced effects different from other early-acting autophagy genes, we next examined an *atg-16.2(ok3224)* mutant and compared it to another early-acting autophagy mutant *atg-4.1(bp501)* for autophagy status and exopher generation. As previously reported in *C. elegans* embryos^30,31^, both mutants displayed increased SQST-1::GFP levels compared to wild-type animals (**Figure 4a**), consistent with a block of autophagy in these animals. Moreover, the two autophagy mutants had pronounced effects on GFP::LGG-1 structures. Specifically, in contrast to the neuronal RNAi (**Figure 2a**), *atg-4.1(bp501)* and *atg-16.2(ok3224)* mutants displayed no reduction of GFP::LGG-1 punctae in nerve-ring neurons, but a marked, and more pronounced increase in lipidation-independent GFP::LGG-1(G116A) (**Figure 4b**) punctae than the neuronal RNAi (**Figure 2b**), indicating that GFP::LGG-1 punctae in the two mutants do not represent functional autophagosomes. To further evaluate autophagy status in these mutants, we performed autophagy flux assays to assess autophagy activity^32^ by injecting them with the late-stage autophagy inhibitor Bafilomycin A (BafA; a compound that blocks acidification of lysosomes^33^). As predicted, BafA did not lead to an increase in GFP-positive punctae in wild-type animals expressing the lipidation-deficient LGG-1(G116A) reporter, consistent with GFP::LGG-1(G116A) punctae not being competent autophagosomes (**Figure S5a**). While BafA injection in wild-type animals increased the number of GFP::LGG-1 punctae, as a reflection of active autophagy, no GFP::LGG-1 punctae increase was observed when BafA was injected into *atg-16.2(ok3224)* and *atg-4.1(bp501)* mutants (**Figure S5b**). Taken together, our results are consistent with neuronal autophagy being blocked in *atg-4.1(bp501)* and *atg-16.2(ok3224)* mutants.

**Figure 4.**
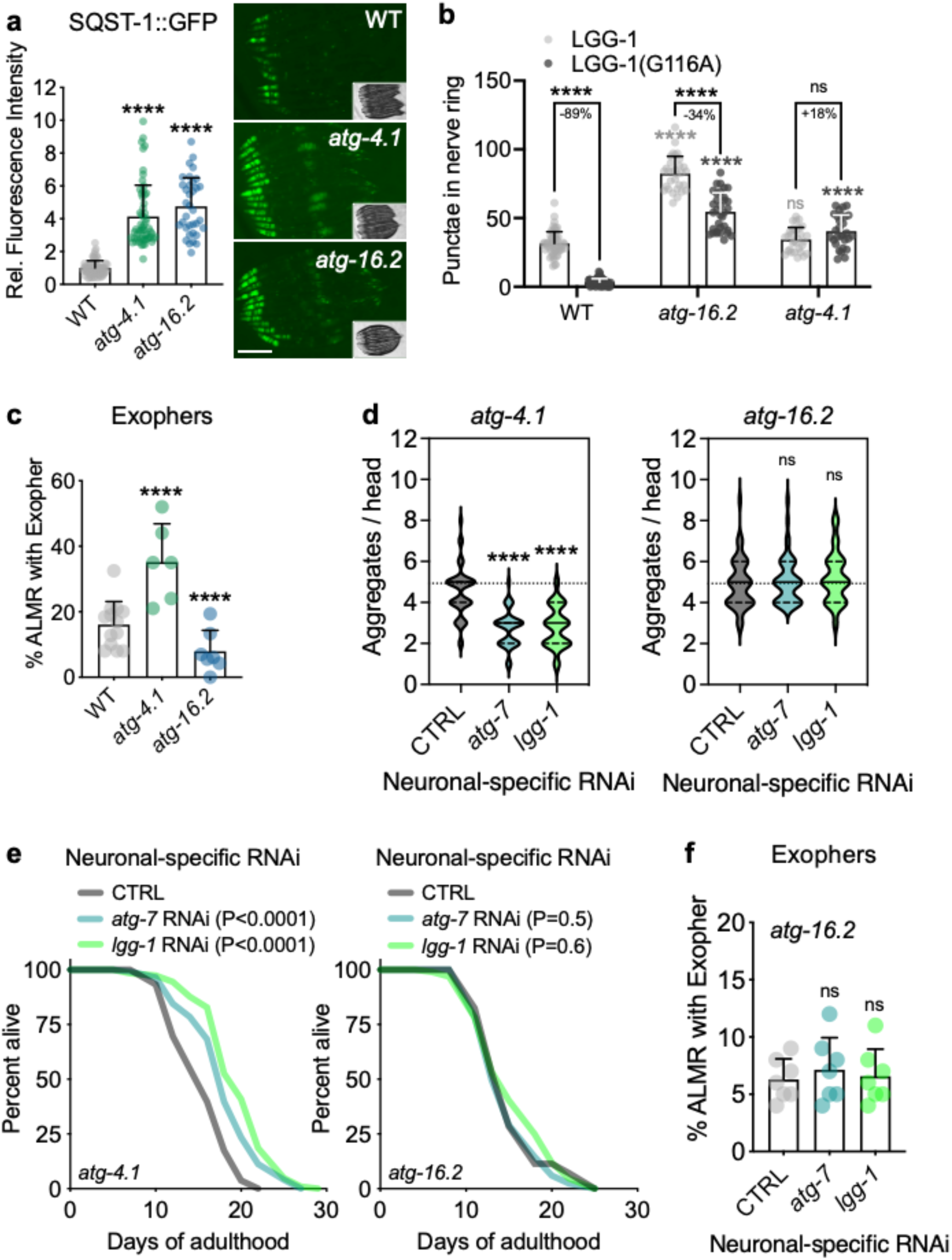
*atg-16.2* is required for exopher formation and for benefits of neuronal inhibition of early autophagy genes independently of degradative autophagy. (**a**)GFP fluorescence intensity in head region in wild-type (WT), *atg-4.1(bp501),* and *atg-16.2 (ok3224)* animals expressing *sqst-1p::sqst-1::gfp.* Data pooled from n=3-5 experiments withN=35-68 animals. Error bars indicate s.d. ****P < 0.0001, by one-way ANOVA. Representative images from one experiment. (**b**) GFP::LGG-1/Atg8- and GFP::LGG-1(G116A)-positive punctae were counted in day 1 wild-type (WT), *atg-4.1(bp501)*, and *atg-16.2(ok3224)* animals. Error bars indicate s.d. of n=3 experiments with N=29-33 animals. ns P > 0.05, by one-way ANOVA. Comparison between strains: ns P > 0.05, ****P < 0.0001, comparison of lipidated and un-lipidated structures: ns P >0.05, ****P < 0.0001, by two-way ANOVA. (**c**) Percent of WT, *atg-4.1(bp501)*, and *atg-16.2(ok3224)* animals expressing *mec-4p::mCherry* with ALMR exophers on day 2 of adulthood. Error bars indicate s.d. of n=6-7 experiments, each with N=28-52 animals. \*\*\*\**P* < 0.0001, by Cochran-Mandel Hansel test. (**d**) Neuronal PolyQ aggregates in *atg-4.1(bp501)* (left) and *atg-16.2(ok3224)* (right) animals capable of neuronal-only RNAi and expressing *rgef-1::Q40::yfp* on day 5 of adulthood fed bacteria expressing empty vector (CTRL), *atg-7*, or *lgg-1/Atg8* dsRNA from hatching on. Violin plots of data pooled from n=3 experiments, with N=45 animals with solid line indicating median and dashed lines indicating quartiles. ns P > 0.05, ****P < 0.0001, by one-way ANOVA. **(e)** Lifespan analyses in *atg-4.1(bp501)* (left) and *atg-16.2(ok3224)* (right) animals capable of neuronal-specific RNAi (*sid-1; rgef-1p::sid-1 + rgef-1p::gfp*) fed bacteria expressing empty vector (CTRL), *atg-7*, or *lgg-1/Atg8* dsRNA from hatching on. Statistical significance determined by log-rank test. See **Table S4** for details and repeats. **(f)** Percent of ALMR neurons with exophers in *atg-16.2(ok3224)* animals capable of neuronal-only RNAi (*sid-1; rgef-1p::sid-1 + rgef-1p::gfp*) and expressing *mec-4p::mCherry* on day 2 of adulthood and fed bacteria expressing dsRNA for empty vector (CTRL), *atg-7*, or *lgg-1/Atg8* from hatching on. Compare with Figure 3b-c, day 2 for wild-type effects. Error bars indicate s.d. of n=7 experiments, each with N=37-62 animals. ns P > 0.05, by Cochran-Mandel Hansel test.

Since whole-body inhibition of autophagy genes *bec-1/Beclin1, atg-7* and *lgg-1/Atg8* increases exopher formation^28,34^, we tested exopher generation on day 2 of adulthood in *atg-4.1* and *atg-16.2* mutants. As expected, we found increased exopher formation in *atg-4.1(bp501)* mutants, but decreased exopher formation in *atg-16.2(ok3224)* mutants (**Figure 4c**), highlighting a novel requirement for *atg-16.2* in exopher biogenesis.

We next tested whether *atg-16.2* could also be required for the benefits of neuronal autophagy-gene inhibition. Specifically, we subjected neuronally RNAi sensitized *atg-4.1(bp501)* and *atg-16.2(ok3224)* mutants to *atg-7(RNAi)* or *lgg-1/Atg8(RNAi)* and found that neuronal-only autophagy-gene inhibition specifically decreased the number of PolyQ aggregates in *atg-4.1(bp501)* mutants, but not in *atg-16.2(ok3224)* mutants (**Figure 4d**). Moreover, neuronal-only autophagy-gene inhibition extended the lifespan of *atg-4.1(bp501)* mutants, but not of *atg-16.2(ok3224)* mutants (**Figure 4e, Table S4**). Consistently, *atg-16.2* was required for the increased exopher formation upon neuronal-only *atg-7(RNAi)* and *lgg-1/Atg8(RNAi);* (**Figure 4f**). The absence of phenotypes in *atg-16.2* mutants was not because these animals were refractory to RNAi, as they responded normally to RNAi clones for genes expressed in muscle, hypodermis, and intestine, or ubiquitously (**Figure S5c**), and neuronal inhibition of the insulin/IGF-1-like receptor *daf-2* increased lifespan in *atg-16.2(ok3224)* mutants (**Table S4**), similar to wild-type animals (**Table S2**). Taken together with our autophagy analyses (**Figures 4a-b, S5a-b**), these results indicate that neuronal-only RNAi of at least *atg-7* and *lgg-1/Atg8* can reduce PolyQ aggregates, and induce exopher formation, as well as extend lifespan, irrespectively of whether canonical autophagy is engaged, consistent with a cargo-degradation independent mechanism in neurons. Our data highlight that this underlying mechanism likely involves *atg-16.2,* which was required for all of the above-mentioned phenotypes.

#### The WD40 domain of ATG-16.2 is dispensable for neuronal autophagy, but required for exopher biogenesis

To further address a mechanistic role for ATG-16.2 in exopher biogenesis, we next tested cell-autonomous functions of ATG-16.2 and performed structure-function analyses by rescuing ATG-16.2 expression in neurons of *atg-16.2* mutants. ATG16 is a conserved protein (**Figure S6a**), containing an ATG5-interacting motif, a coiled-coil domain, and a seven bladed β-propeller WD40 domain, mediating protein interactions at its C terminus (**Figure 5a-b**). While the *S. cerevisiae* Atg16 orthologue does not contain a WD40 domain (**Figure S6a**), the WD40 domain has been shown to be dispensable for canonical, degradative autophagy functions in metazoans^13^. To test the function of the *C. elegans* ATG-16.2 WD40 domain, we created and analyzed an ATG-16.2ΔWD40 protein (**Figure 5a**), and an ATG-16.2(F394A) protein carrying a point mutation of phenylalanine 394 to alanine. This phenylalanine is conserved (**Figure S6a**) and important for the WD40 function in ATG16L1 in mammalian cells^13^. We expressed each of these proteins, as well as a full-length ATG-16.2 from the *rgef-1* promoter in neurons of *atg-16.2(ok3224)* mutants and found their gene expression levels to be similar to wild-type levels (**Figure S6b**). Since *atg-16.2(ok3224)* mutants displayed an increased number of GFP::LGG-1 punctae (**Figure 4b, S5b**) due to blocked autophagy (**Figure 4a**), we first monitored GFP::LGG-1 punctae in the neurons in *atg-16.2(ok3224)* and found that neuronal rescue of full-length ATG-16.2, ATG-16.2ΔWD40, and ATG-16.2(F394A) all similarly returned the number of neuronal GFP::LGG-1 punctae compared to WT levels (**Figure 5C**). These results indicate that the ATG-16.2 constructs are functional, and that the ATG-16.2 WD40 domain is not required for neuronal autophagy in adult *C. elegans*, consistent with previous studies in *C. elegans* embryos31 and mammalian cells^13^.

**Figure 5.**
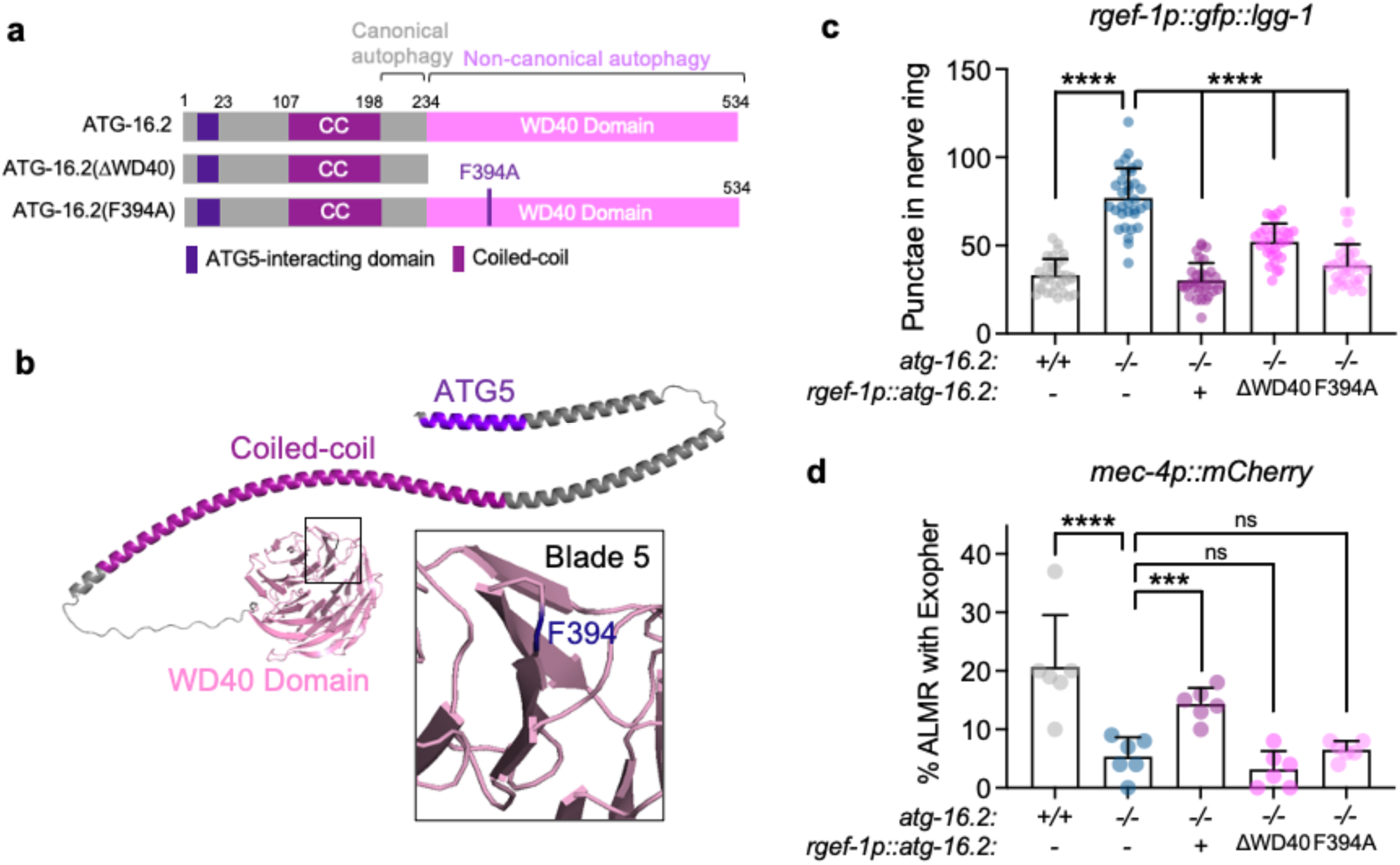
The WD40 domain of ATG-16.2 is dispensable for autophagosome formation but required for exophergenesis. **(a)** Schematic of ATG-16.2 rescue constructs. Full-length ATG-16.2 protein includes ATG5-interacting motif, coiled-coil domain (CC), and WD40 domain. ATG-16.2ΔWD40 is a truncated ATG-16.2 protein containing amino acids 1-234. ATG-16.2(F394A) protein contains a point mutation of phenylalanine to alanine in position 349 in the WD40 domain. See **Figure S6A** for primary structure of ATG-16.2. (**b**) AlphaFold model of ATG-16.2 (UniProt Q09406). The N-terminal region of ATG-16.2 is predicted to contain a helical structure with a conserved ATG5-interacting motif and a CC domain. The C terminus contains a seven-bladed WD40 beta-propeller domain. F394 is located on blade 5 on the linker between β-sheets B-C (enlarged). (**c**) Neuronal GFP::LGG-1 punctae were scored in wild-type (WT), *atg-16.2(ok3224)*, and *atg-16.2(ok3224)* mutants transgenically expressing either full-length *atg-16.2*, *atg-16.2* lacking the WD40 domain (ΔWD40), or *atg-16.2(F394A)* (F394A) from the neuronal *rgef-1* promoter. Error bars indicate s.d. of n=3 experiments, with N=28-29 animals. ****P < 0.0001, by one-way ANOVA. (**d**) Exophers were scored in wild-type (WT), *atg-16.2(ok3224)*, and *atg-16.2(ok3224)* mutants expressing either full-length *atg-16.2, atg-16.2* lacking the WD40 domain (ΔWD40), or *atg-394A)* (F394A) from the neuronal *rgef-1* promoter. Quantification of percent of ALMR neurons with exophers on day 2 of adulthood. Error bars indicate s.d. of n=6 experiments, each=23-58 animals. ns P > 0.05, *** P <0.001, ****P < 0.0001 by Cochran-Mandel Hansel test.

Since *atg-16.2* was required for exopher biogenesis (**Figure 4c**), we next tested whether *atg-16.2* and its WD40 domain were cell-autonomously required for exopher formation. We found that *atg-16.2(ok3224)* mutants expressing full-length ATG-16.2, but neither of the WD40 domain-defective ATG-16.2 constructs (ΔWD40 and F394A), significantly increased exopher numbers to near wild-type levels (**Figure 5d**). These results indicate that a neuronal function of ATG-16.2 involving its WD40 domain is important for exopher formation.

### The WD40 domain of ATG-16.2 is required for the benefits of neuronal inhibition of autophagy genes *atg-7* and *lgg-1/Atg8*

Lastly, we analyzed if neuronal ATG-16.2 was involved in the effects mediated by neuronal inhibition of early-acting autophagy genes. To this end, we created *atg-16.2(ok3224)* mutants capable of neuronal RNAi and expressing full-length neuronally expressed ATG-16.2, or WD40 domain-deficient ATG-16.2ΔWD40 and ATG-16.2(F394A). We subjected these strains to neuronal RNAi of *atg-7* or *lgg-1/Atg8* and examined them for neuronal PolyQ aggregates (**Figure 6a-c**), exopher production (**Figure 6d-f**), and lifespan (**Figure 6g-i**, **Table S5**). In the animals rescued with full-length ATG-16.2, neuronal knockdown of *atg-7* or *lgg-1/Atg8* reduced neuronal PolyQ (**Figure 6a**), increased exopher production (**Figure 6d**), and extended lifespan (**Figure 6g**), similar to wild-type animals. In contrast, neuronal expression of ATG-16.2ΔWD40 or ATG-16.2(F394A) failed to rescue the phenotypes of neuronal PolyQ reduction (**Figure 6b-c),** exopher genesis (**Figure 6e-f**), and lifespan extension (**Figure 6h-i**). These results demonstrate that the WD40 domain of ATG-16.2 is autonomously required for neuronal PolyQ reduction, increased exopher production, and lifespan extension in *atg-16.2(ok3224)* mutants following neuronal knockdown of early-acting autophagy genes.

**Figure 6.**
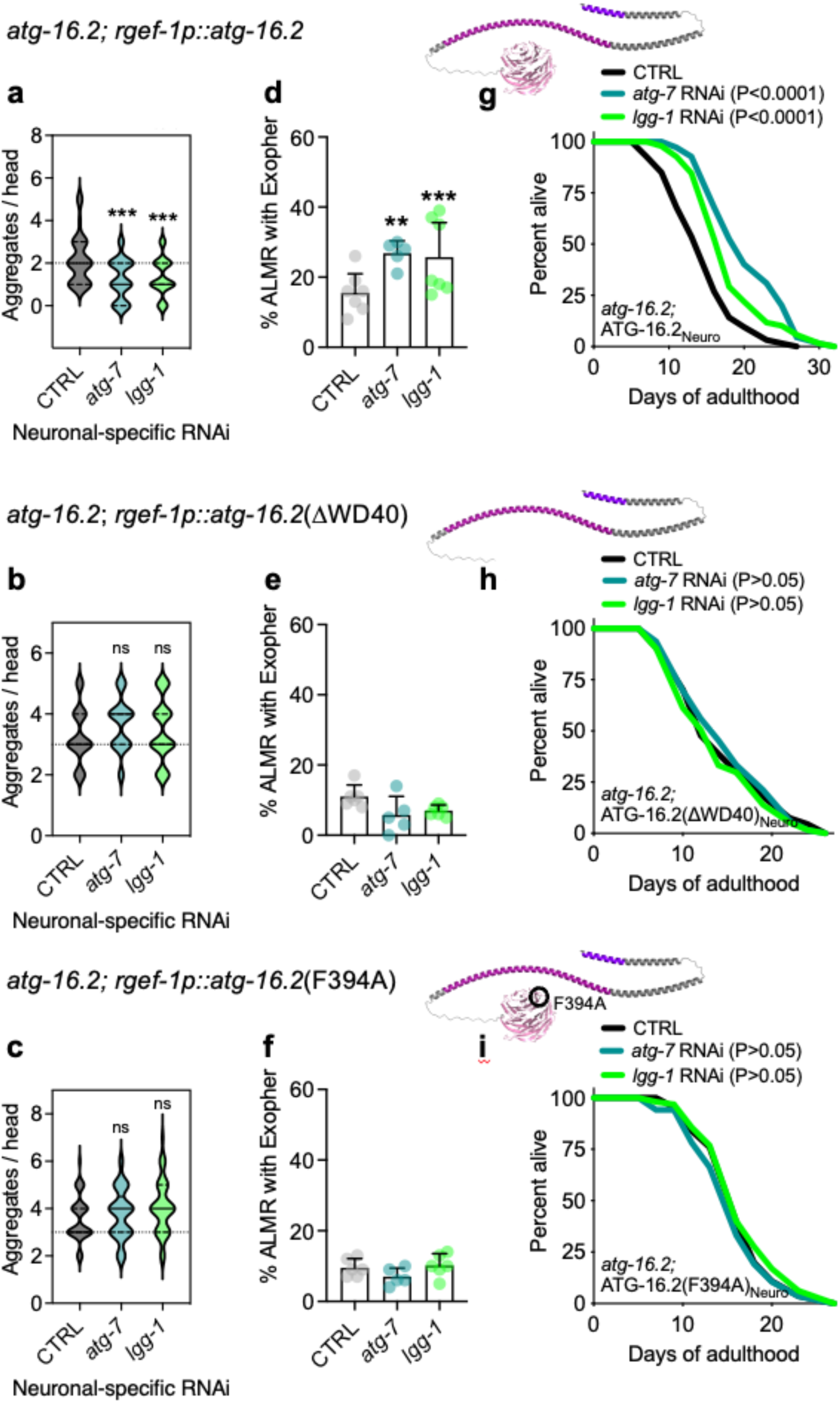
The WD40 domain of ATG-16.2 is required for the reduced PolyQ aggregates, the increased exopher percentage, and the lifespan extension observed upon neuronal down of early-acting autophagy genes. **(a-c)** Neuronal PolyQ aggregates in *atg-16.2(ok3224)* animals expressing *rgef-1::Q40::yfp* on day 5 of adulthood fed bacteria expressing empty vector (CTRL), *atg-7*, or *lgg-1/Atg8* dsRNA from hatching on. Full-length *atg-16.2* **(a),** *atg-16.2ΔWD40* **(b),** or *atg-16.2(F394A)* **(c)** were expressed from the pan-neuronal promotor *rgef-1 in atg-16.2(ok3224)* mutants. Violin plots of data from three independent experiments, each with N=10 animals with solid line indicating median and dashed lines indicating quartiles. ns P > 0.05, ***P < 0.001, by one-way ANOVA. (**d-f**) Exophers were scored in day 2 *atg-16.2(ok3224)* mutants expressing *mec-4p::mCherry* and fed bacteria expressing CTRL or *atg-7* or *lgg-1/Atg8* dsRNA from hatching. Complementation was tested in *atg-16.2(ok3224)* mutants by expressing full-length *atg-16.2* **(d),** *atg-16.2ΔWD40* **(e),** or *atg-16.2(F394A)* **(f)**, from the pan-neuronal promotor *rgef-1*. Quantification of percent of ALMR neurons with exophers. Error bars indicate s.d. of n=5-7 experiments, each with N=28-54 animals. ns P > 0.05, ** P <0.01, ***P < 0.001 by Cochran-Mandel Hansel test. **(g-i)** Lifespan analysis of *atg-16.2(ok3224)* mutants capable of neuronal-only RNAi and fed bacteria expressing CTRL, *atg-7*, or *lgg-1/Atg8* dsRNA from hatching on. *atg-16.2(ok3224)* mutants were rescued by expressing full-length *atg-16.2* **(g),** *atg-16.2ΔWD40* **(h),** or *atg-16.2(F394A)* **(i)** from the pan-neuronal promotor *rgef-1*. Statistical significance determined by log-rank test. See Supplementary **Table S2 and Table S5** for details and repeats.

Collectively, our results show a novel role for the WD40 domain of neuronal ATG-16.2 in exopher biogenesis and provide a potential mechanism by which pan-neuronal inhibition of early-acting autophagy genes leads to decreased PolyQ aggregation, possibly via increased exopher formation in at least some neurons. Finally, our results highlight a novel role for neuronal ATG-16.2 in lifespan extension induced by early autophagy-gene inhibition.

## DISCUSSION

Here we demonstrated that RNAi of early-acting autophagy genes in *C. elegans* neurons significantly extended lifespan, autonomously protected neuron morphology in aged animals, and improved neuronal proteostasis by reducing neuronal PolyQ aggregates and increasing the secretion of exophers, which are large extracellular vesicles. These effects were likely independent of canonical autophagy, i.e., lysosomal degradation, since downregulation of late-acting autophagy genes did not induce the same benefits, all neuronal-only autophagy gene RNAi treatments, as expected, induced an autophagy block, and because *atg-4.1* mutants, which have defects in autophagy, displayed the same phenotypes as wild-type animals following downregulation of early-acting autophagy genes. Instead, we found that the extension of lifespan and improvement of neuronal proteostasis as well as increased exopher biogenesis after inhibition of early-acting autophagy genes in neurons required ATG-16.2, and particularly the WD40 domain-related function of ATG-16.2. Our data uncover a role for the ATG-16.2 WD40 domain in the formation of exophers, which have been shown to secrete cytosolic material, including PolyQ aggregates from *C. elegans* neurons, especially upon disruption of proteostatic pathways28. We thus propose a model in which the inhibition of early-acting autophagy genes leads to increased exopher formation in a manner dependent on the WD40 domain of ATG-16.2; this can explain improved neuronal proteostasis and may contribute to lifespan extension in a direct or indirect fashion.

We here identified several new functions for the autophagy protein ATG-16.2 in adult *C. elegans.* Mammalian ortholog ATG16L1 mainly functions in the ATG12-ATG5/ATG16 complex to mediate Atg8 conjugation to double-membrane autophagosomes, but can also via its highly conserved WD40 domain facilitate association of ATG8/LC3 to non-autophagosomal membranes, including endolysosomal membranes or single-membrane vesicles involved in LC3-associated phagocytosis (LAP) in mouse macrophages^13^. We here found the WD40 domain of *C. elegans* ATG-16.2 to be required for basal exopher formation as well as the increase in exopher formation observed following neuronal inhibition of early-acting autophagy genes *atg-7* and *lgg-1/Atg8*. In addition to *C. elegans*^35,36^, exophers have been observed in mammalian cardiomyocytes^36^ and neurons^35^, but exopher biogenesis is not well understood molecularly. Our observations raise the interesting question of whether *C. elegans* ATG-16.2, via its WD40 domain, can get recruited to exopher membranes, and whether such potential non-canonical functions of ATG-16.2 would involve the recruitment and conjugation of ATG8 proteins to exopher membranes via its association with the ATG-12-ATG-5 complex. To this end, neuronal-only reduction of *lgg-1/Atg8* not only increased exopher formation, but also improved neuronal proteostasis, and increased lifespan similar to other early-acting autophagy genes. While this could point towards an ATG8-independent function of ATG-16.2, the partial knockdown of neuronal *lgg-1*/Atg8 (∼40% remaining after the RNAi knockdown treatment) could be sufficient for non-canonical, but insufficient for canonical autophagy. Alternatively, ATG-16.2 could have an autophagy-independent function, i.e., independent of its role in ATG8 lipidation, similar to mammalian ATG16L1, which can facilitate hormone secretion via dense-core vesicles^37^ and exosome production^38^ in cellular models, as well as neuropeptide production in *Drosophila*^39^ in an autophagy-independent manner. Future experiments are needed to address the mechanism by which ATG-16.2 regulates exopher formation in response to neuronal-only autophagy gene knockdown.

Neuronal autophagy is important for protein homeostasis in the nervous system, since conditional knockdown of autophagy genes *Atg5* and *Atg7* in the central nervous system of mice leads to neurodegenerative phenotypes^40,41^, and loss-of-function mutations in the endo-lysosomal pathway frequently cause neurological disorders^42,43^. Autophagy has been shown to degrade aggregating proteins, including proteins with PolyQ stretches^44^. In line with these observations, non-neuronal inhibition of all autophagy-related genes tested in *C. elegans* increased neuronal PolyQ formation (this study and^17-19^). Surprisingly, and in contrast to the non-neuronal findings, we here report that neuronal-only inhibition of early-acting autophagy genes reduced PolyQ aggregate load in neurons. These observations raise several questions, including how can neuronal inhibition of early-autophagy genes improve PolyQ aggregation and proteostasis in neurons, and what is the fate of neuronal PolyQ aggregates upon inhibition of early-acting autophagy genes in neurons? While these important questions remain to be experimentally investigated, we favor a model where secretion of aggregating proteins from neurons, could be induced in response to neuronal inhibition of early-acting autophagy genes likely via extruded exophers, which can contain aggregated proteins28. This would be consistent with recent studies that have shown that PolyQ proteins and aggregates can be transmitted from cell-to-cell in *C. elegans*^45^, *Drosophila*^46^, as well as in induced pluripotent stem cells derived from Huntington’s disease patients and transplanted into mouse brains^47^, possibly via extracellular vesicles^48,49^. Moreover, inhibition of autophagosome formation by silencing of early-acting autophagy genes *Atg5* or *Vps34* in different neuronal cell culture models promotes exosomal secretion of aggregate-prone proteins, such as α-synuclein^50^, prions^51^, or an amyloid precursor^52^. It will be interesting to investigate if these secretory events require ATG16 proteins, and their WD40 domain. Likewise, it will be important to better understand the physiological role of potentially secreted, aggregated proteins in cell communication, noting that our studies demonstrated a direct correlation between decreased neuronal aggregate load and increased lifespan.

An especially surprising observation made in this study was that neuronal-only inhibition of early-acting autophagy genes from hatching or from day 1 of adulthood, using the same RNAi bacterial clones that shorten the lifespan of several long-lived *C. elegans* mutants (which would experience gene inhibition in non-neuronal tissues)1 caused normal *C. elegans* to live ∼20-30% longer [we note that a recent *C. elegans* study reported that neuronal-only inhibition of *bec-1/Beclin1* from day 1, using a different RNAi-permissible strain^16^, reduced lifespan, while *bec-1/Beclin1* and *vps-34* neuronal-only knockdown from day 9 instead resulted in lifespan extension^26^; we do not know the reason for the discrepancy in results during neuronal-only day 1 *bec-1/Beclin1* inhibition besides use of different RNAi strains]. Our lifespan results are notable since autophagy gene Rubcn (Run domain Beclin-1 interacting and cysteine-rich containing protein) negatively regulates autophagy, and whole-life, neuronal-only RNAi of *rub-1/Rubcn* in *C. elegans* extends lifespan via increasing autophagy flux^53^. Thus, canonical autophagy induction in neurons can induce beneficial effects on neuronal proteostasis and lifespan in *C. elegans*. Our data, on the other hand, suggest that the inhibition of early-autophagy autophagy genes triggers a longevity mechanism independent of degradative autophagy, and mediated by the WD40 domain of ATG-16.2. Considering the requirement of this highly conserved domain in non-canonical autophagy gene functions in mammals, we propose a similar role in adult *C. elegans* neurons with possible secretion of lifespan-extending signals, either in exophers or different types of *atg-16.2*-dependent secretory vesicles. In future experiments, it will be interesting to address if and how the neuronal secretome may be affected in *C. elegans atg-16.2* mutants.

In sum, we here demonstrated that the inhibition of early-acting autophagy genes in neurons extended *C. elegans* lifespan, improved neuronal proteostasis, and increased exopher formation mediated by the autonomous, WD40 domain-related function of ATG-16.2. If this mechanism is conserved in other organisms, modulating neuronal autophagy genes may provide a method to improve lifespan and neuronal proteostasis. In addition, as the WD40 domain of *C. elegans* ATG-16.2 was critical for lifespan determination, we speculate that the non-canonical ATG16 functions may play a broader role in organismal aging.

## AUTHOR CONTRIBUTIONS

C.K., Y.Y., and M.H. designed the experiments. Y.Y., C.K., M.L.A., T.M.M., and M.H. constructed strains. Y.Y., C.K., E.H.C., C.M.L., K.P., M.B., L.H.S., T.M.M., A.S., and M.H. performed lifespan experiments. Y.Y. and C.K. counted PolyQ aggregates. Y.Y. and E.H.C. conducted healthspan assays. Y.Y., C.M.L. and C.K. measured RNAi efficiency. Y.Y. conducted neuronal branching and Dye-filling experiments and cloning. C.K. and M.L.A. monitored exopher formation. C.K. performed all autophagy measurements and qRT-PCR. Y.Y., C.K., and M.H. wrote the manuscript, with input from coauthors.

## COMPETING INTEREST

The authors declare no competing interests.

## ACKNOWLEDGEMENTS

We thank members of the Hansen and Kumsta labs for comments on the manuscript. We thank Dr. Xingyu She, Dr. Ee Phie Tan, and Shaun Lim from the Hansen lab for creating some of the strains used in this study. We thank Dr. Hong Zhang for sharing construct *atg-16.2p::atg-16.2::gfp* and Dr. Barth Grant for sharing the hygromycin resistance-containing plasmid. We thank Dr. Oliver Florey for suggestions on making the *rgef-1p::atg-16.2(F394A)* construct. We thank Drs. Nina Riehs and Cynthia Kenyon for sharing unpublished data on neuronal-only RNAi of autophagy genes ahead of publication. Some of the nematode strains used in this work were provided by the Caenorhabditis Genetics Center (University of Minnesota), which is supported by the NIH–Office of Research Infrastructure Program (P40 OD010440). We acknowledge the *C. elegans* Gene Knockout Project at the Oklahoma Medical Research Foundation, which was part of the International *C. elegans* Gene Knockout Consortium, for generating *atg-16.2(ok3224)* mutation. This work was funded by NIH/NIA grants R37AG056510 to MD, F31AG066405 to MLA, 664 and AG039756 and a Breakthrough in Gerontology Award from American Association ng Research to MH.

## METHODS

### *C. elegans* strains and maintenance

*C. elegans* strains were maintained and cultured at 20°C using *Escherichia coli* OP50 as a food source54 unless RNAi was initiated. For RNAi experiments, animals were grown on HT115 bacteria from the time of RNAi initiation (see below). See **Table S6** for all strains used and created for this study. For maintenance and selection of animals expressing hygromycin resistance (HygR) marker, animals were grown on 6 cm NGM plates that were supplemented with 250 μl of 5 mg/ml of hygromycin B (GoldBio) in M9.

### Construction of transgenic strains

The *rgef-1p::sid-1* vector was made by Gateway cloning. The *rgef-1* pan-neuronal promoter, *sid-1* open reading frame, and the *unc-54* 3’-UTR were cloned into the Gateway destination vector pDEST_R3R4. The final product *rgef-1p::sid-1* (pMH1141) was verified by sequencing. To construct the *rgef-1p::atg-16.2* vector, cDNA of *atg-16.2* was amplified by PCR from vector *atg-16.2p::atg-16.2::gfp*31 and ligated with pMH1307 (pSM vector containing *rgef-1* promotor, between NheI and SalI) by Gibson assembly. The final product *rgef-1p::atg-16.2* (pMH1387) was verified by sequencing. The construct *rgef-1p::atg-16.2(F394A)* was created by site-directed mutagenesis, in which the coding sequence of phenylalanine 394 (TTT) was changed to alanine 394 (GCT) in construct pMH1387. The final product *rgef-1p::atg-16.2(F394A)* (pMH1388) was verified by sequencing. To construct the *rgef-1p::atg-16.2ΔWD40* vector, cDNA of *atg-16.2delC,* which encodes ATG-16.2(1-223), was amplified by PCR from vector *atg-16.2p::atg-16.2*31 with added stop codon and ligated with pMH1307 (between Nhe I and Sal I) by Gibson assembly. The final product *rgef-1p::atg-16.2ΔWD40* (pMH1389) was verified by sequencing. Plasmid DNA was prepared using the Mini Prep kit (Qiagen). Transgenic animals expressing an extrachromosomal array were created by gonadal microinjection of plasmids of interest with indicated co-injection marker into indicated strains in **Table S6**. A list of plasmids used and made in this study is provided in **Table S7**. Primer information for plasmid construction is available in **Table S8**.

### RNA interference

RNA interference (RNAi) was performed by feeding *C. elegans* with bacteria expressing double-stranded RNA (dsRNA) against the gene of interest. RNAi clones used in this study listed in **Table S7** and obtained from the Ahringer3 or the Vidal55 RNAi libraries. All RNAi clones were verified by sequencing. Two empty vector controls were used; the original L4440 plasmid, and L4440, digested with Eco RV and religated, which eliminated 114 bp (Eco RV restriction site was used to insert dsRNA) (pMH1355). For RNAi experiments, HT115 bacteria were grown in liquid LB medium containing 0.1 mg/ml carbenicillin (Bio Pioneer) and 80 µl aliquots of bacteria were spotted onto 6 cm NGM plates with carbenicillin. Bacteria were allowed to grow for 1-2 days at room temperature. Before use, 80 µl 0.1 M IPTG (Promega) were added to the bacterial lawn to induce dsRNA expression before eggs (whole-life RNAi), larvae (L1 or L4 stage), or adults (adult-only RNAi) were transferred to RNAi plates. Neuronal RNAi was performed in *sid-1(qt9)* mutants with *sid-1* rescue under control of the pan-neuronal promoter *rgef-1p*. RNAi efficiency in mutants and neuronal RNAi strains was assessed with whole-life RNAi treatment on day 2 of adulthood. RNAi specific against genes with muscle function: *unc-112* and *unc-22* RNAi penetrance were measured by paralysis, and twitch-like movements, respectively. RNAi specific against genes with hypodermal function: *tsp-15*, *bli-1*, and *bli-3* RNAi penetrance was determined by the appearance of blisters, whereas *die-1* RNAi led to larval arrest and crippled appearance. RNAi specific against genes with intestinal function: *elt-2* and *pept-1* RNAi penetrance was determined by a small phenotype and a clear and restricted appearing intestine. RNAi against the ubiquitously expressed *rpl-2* led to larval arrest. RNAi specific against genes with neuronal function: *snb-1* and *unc-13* RNAi caused shrinker phenotype. Statistical analysis was done using one-way ANOVA using GraphPad Prism.

### Lifespan analysis

Lifespan was measured at 20°C as previously described^56^. Briefly, synchronized animals were transferred onto 6 cm NGM plates seeded with *E. coli* OP50. Six plates were used for each strain with 20 animals per plate. The L4 larval stage was recorded as day 0 of lifespan, and animals were transferred every other day to a new NGM plate throughout the reproductive period. For lifespan experiments on RNAi bacteria, animals were fed dsRNA-expressing or control bacteria from hatching (whole-life), from larvae (L1 or L4 stage), or as adults (adult-only RNAi). Animals were scored as dead if they failed to respond to gentle prodding with a platinum wire pick. Censoring occurred if animals desiccated on the edge of the plate, escaped, ruptured, or suffered from internal hatching. Statistical analysis was performed using STATA software (Stata Corp) or Oasis.2^57^. *P*-values were calculated using the log-rank (Mantel–Cox) method. All lifespan experiments performed for this study are numbered to indicate which RNAi clones were tested together in the same experiment, and listed in **Tables S1-S5.**

### PolyQ aggregation

The number of neuronal PolyQ aggregates was counted in animals expressing *rgef-1p::Q40::yfp* in the nerve-ring of individual animals on day 5 of adulthood^17,18^. Animals were raised at 20°C on control HT115 bacteria or RNAi bacteria against the gene of interest. Animals were lined up on 6 cm NGM plates without food after anaesthetization with M9 medium containing 0.1% NaN_3_. Neuronal PolyQ aggregates were counted under a Leica DFC310 FX camera at 20x magnification.

### Neurite branching analysis

Neurite branches were counted as previously described^22^. Briefly, eggs were raised on control HT115 bacteria or RNAi bacteria plates for whole-life RNAi. Branches from ALM and PLM mechanosensory neurons were counted on day 15 of adulthood. A branch was scored when a visible GFP-labeled branch was observed emanating from ALM or PLM mechanosensory neurons visualized using a Zeiss Imager Z1 including apotome at 63x magnification. 20 animals were scored for each condition and results were analyzed using Cochran-Mandel Hansel analysis in Microsoft Excel.

### Exopher measurements

Exophers were counted as previously described 29. Briefly, animals expressing *mec-4p::mCherry* were raised on OP50 or HT115 bacteria expressing empty vector or dsRNA from hatching on. Animals were scored on day 1-3, or just on day 2 of adulthood for exopher occurrence in a binary manner. Animals were either scored live on a Kramer dissecting scope with a 20x objective or mounted on a microscope slide in a drop of M9 media on an agarose pad, both containing 0.1% NaN_3_ as anesthesia using a Zeiss Imager Z1 including apotome at 25x magnification. Each trial was graphed as a percentage of ALMR exophers. We used Cochran-Mandel Hansel analysis for *P*-value calculation of at least 5 or more biological trials. For most exophers measurements, the scoring was performed blinded.

### Fluorescence intensity measurements

Fluorescence intensity of GFP in nerve-ring neurons was measured in animals capable of neuronal RNAi to determine neuronal RNAi knockdown efficiency after feeding animals bacteria carrying empty vector control or expressing dsRNA against *gfp*. Fluorescence intensity of SQST-1::GFP was measured in either the head region or whole body on the indicated days of adulthood either raised on OP50 bacteria or after animals were subjected to the indicated neuronal RNAi treatment. At least three independent experiments were performed per strain. Animals were imaged on empty NGM plates after anaesthetization with M9 medium containing 0.1% NaN_3_. Images were acquired with a Leica DFC310 FX camera at 400 ms exposure. The mean fluorescence intensity was measured in the nerve-ring neurons by outlining the same sized circular region of interest using FIJI software (National Institutes of Health) per animal and in whole body by outlining each worm, normalized to day 1 of adulthood or control RNAi. Data was combined and analyzed using appropriate statistical test (see figure legends) using GraphPad Prism.

### Healthspan measurements

Thrashing ability (i.e., swimming), pharyngeal pumping, and progeny production were assayed as measures of healthspan. For thrashing assays, animals on the indicated days of adulthood were transferred onto a 6 cm NGM media plate containing a drop of M9 medium, and body bends of 14-20 animals were counted for 20 s on a Leica stereoscope. Pharyngeal pumping was measured on the indicated days of adulthood by counting the grinder movements in the terminal pharyngeal bulb of 14–20 animals for 30 s on a Leica stereoscope. For the assessment of a progeny production, 10 animals were singled on 6 cm NGM plates at the L4 larval stage and transferred daily onto fresh plates during the self-fertile reproductive span. The number of eggs/larvae produced by each animal per day was counted. Data were analyzed by appropriate statistical test (see figure legends) using GraphPad Prism.

### Dye-filling assay

Dye-filling experiment were performed on day-5 adulthood. Animals were raised at 20°C on corresponding RNAi plates for whole-life RNAi. On the day of experiment, 20 worms were transferred into 150 μl of a 10 ng μl-1 DiI (Invitrogen) solution diluted in M9 buffer. Animals were incubated for 2 hours at 20°C and then transferred to a fresh NGM plate seeded with OP50 bacteria to let them crawl for 1 hour. Animals were mounted on a 2% agarose pad in M9 medium containing 0.1% NaN_3_ and imaged using a Zeiss Imager Z1 including apotome.2 with a Hamamatsu orca flash 4LT camera and Zen 2.3 software. 10 animals were scored for each condition in three independent experiments.

### Autophagy measurements

GFP::LGG-1 and lipidation-deficient GFP::LGG-1(G116A) punctae were counted in the nerve-ring neurons of animals expressing *rgef-1p::gfp::lgg-125* and *rgef-1p::gfp::lgg-1(G116A)*18, respectively. For all experiments, animals were raised at 20°C and assessed for autophagy status on day 1 of adulthood. Autophagy flux assays were performed as previously described 32. Briefly, bafilomycin A1 (BafA) (BioViotica) or vehicle (dimethylsulfoxide, DMSO) (Sigma) was injected into *rgef-1p::gfp::lgg-1-* or *rgef-1p::gfp::lgg-1(G116A)*-expressing animals. BafA was resuspended in DMSO to a stock concentration of 25 μM. The co-injection dye Texas Red Dextran 3000MW (Molecular Probes) was resuspended in water to a stock concentration of 25 mg ml−1. The BafA or DMSO injection solution was injected close to the terminal pharyngeal bulb and animals were allowed to recover on 6 cm NGM plates with OP50 for 2 h. Surviving animals with intact nerve rings that scored positive for the red dye were used for GFP::LGG-1 punctae quantification. For imaging and punctae quantification, animals were mounted on a 2% agarose pad in M9 medium containing 0.1% NaN_3_ and GFP-positive punctae were counted using a Zeiss Imager Z1 including apotome.2 with a Hamamatsu orca flash 4LT camera and Zen 2.3 software. Z-stack images were acquired at 1.0 µm slice intervals. 6–12 animals were imaged for each condition and results were combined from three independent experiments and analyzed using one-way analysis of variance (ANOVA) using GraphPad Prism. To assess function of *atg-16.2* constructs for rescuing neuronal autophagosome phenotype in *atg-16.2* mutant, pictures were imaged and analyzed double-blinded.

### Quantitative RT-PCR

Quantitative reverse transcriptase–PCR was performed as previously described18. Briefly, total RNA was isolated from a synchronized population of ∼2,000 day 1 animals raised on OP50 bacteria on 6 cm NGM plates, supplemented with hygromycin B where applicable. After harvesting, the animals were flash frozen in liquid nitrogen. RNA was extracted with TRIzol (Life Technologies), purified using a Qiagen RNeasy kit, and subjected to an additional DNA digestion step (Qiagen DNase I kit). Reverse transcription (1 μg RNA per sample) was performed using M-MuLV reverse transcriptase (Roche) and random 9-mer primers (New England Biolabs). Quantitative reverse transcriptase–PCR was performed using SYBR Green Master Mix (Roche) in a CFX384 machine (BioRad). A standard curve was obtained for each primer set by serially diluting a mixture of different complementary DNAs and the standard curves were used to convert the observed CT values to relative values. Three biological samples were analyzed, each with three technical replicates. The average and s.e.m. were calculated for each mRNA. mRNA levels of target genes were normalized to the mean of the following housekeeping genes: *nhr-23* (nuclear hormone receptor), *pmp-3* (putative ABC transporter), and *cyn-1* (cyclophilin). Primer sequences are listed in **Table S8**. Data are displayed as relative values compared with controls. Data were analyzed using one-way ANOVA (GraphPad Prism).

## SUPPLEMENTAL FIGURE LEGENDS

**Figure S1.**
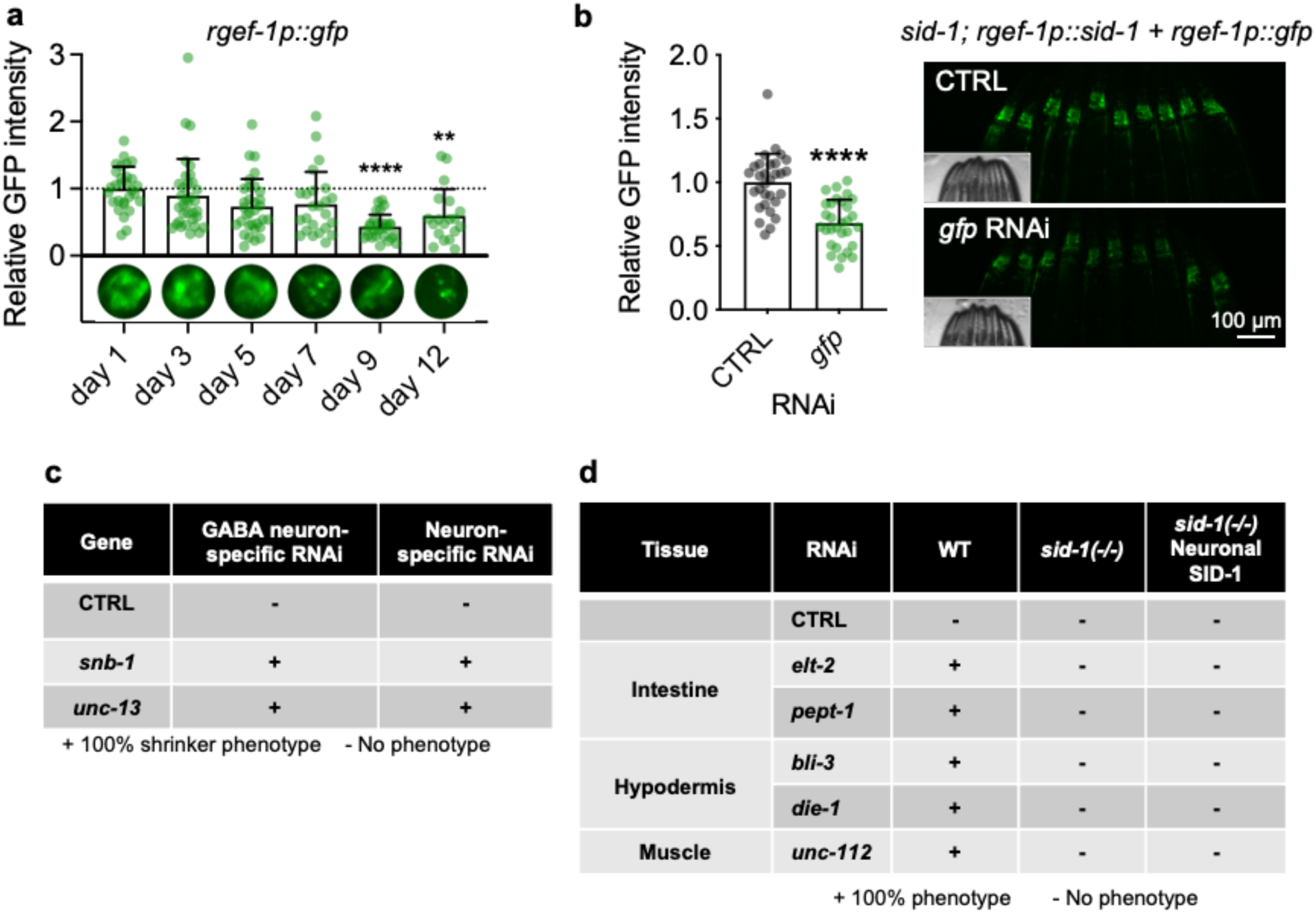
Neuronal expression of SID-1, an RNA channel protein, leads to RNAi-competent neurons without affecting neuronal proteostasis. **(a)** Analysis of relative GFP levels in the nerve ring of animals expressing *rgef-1p::gfp* on day 1 to day 12 of adulthood, normalized to day 1. Error bars indicate s.d. of n=3 experiments, with N=19-34 animals. ***P < 0.001, ****P < 0.0001, by one-way ANOVA. **(B)** Analysis of relative GFP level in the nerve ring in animals capable of neuronal-only RNAi (*sid-1; rgef-1p::sid-1 + rgef-1p::gfp*) fed bacteria expressing empty vector (CTRL) or *gfp* dsRNA from hatching and analyzed on day 1 of adulthood. Error bars indicate s.d. of n=3 experiments with N=30 animals. ****P < 0.0001, by Student’s t-test. **(C)** Analysis of shrinker phenotype in day 1 animals capable of neuronal-only RNAi (*sid-1; rgef-1p::sid-1 + rgef-1p::gfp*) or capable of GABA neuron-specific RNAi (*rde-1; unc-47p::rde-1::SL2::sid-1*). Animals were subjected to bacteria expressing empty vector (CTRL), *unc-13*, or *snb-1* dsRNA whole-life for two generations. Three replicates for each gene with N=10 animals in each repeat. +: 100% of animals showed shrinker phenotype when poked at the head region with a platinum wire. -: No phenotype observed when poked at the head region with a platinum wire. **(D)** Wild-type animals (N2, WT), RNAi-resistant animals (*sid-1* mutants), and animals capable of neuronal-only RNAi (*sid-1; rgef-1p::sid-1 + rgef-1p::gfp*) were fed bacteria from hatching expressing empty vector (CTRL), or dsRNA against a specific gene in different tissues, and phenotypes were analyzed on day 1 of adulthood. Phenotypes for knockdown of each gene: *elt-2*: small and clear, *pept-1*: lethal, *bli-3*: blisters, *die-1*: lethal, *unc-122*: paralysis. Three replicate for each gene with N=20 animals in each repeat. +: 100% of animals showed phenotype. -: No phenotype observed.

**Figure S2.**
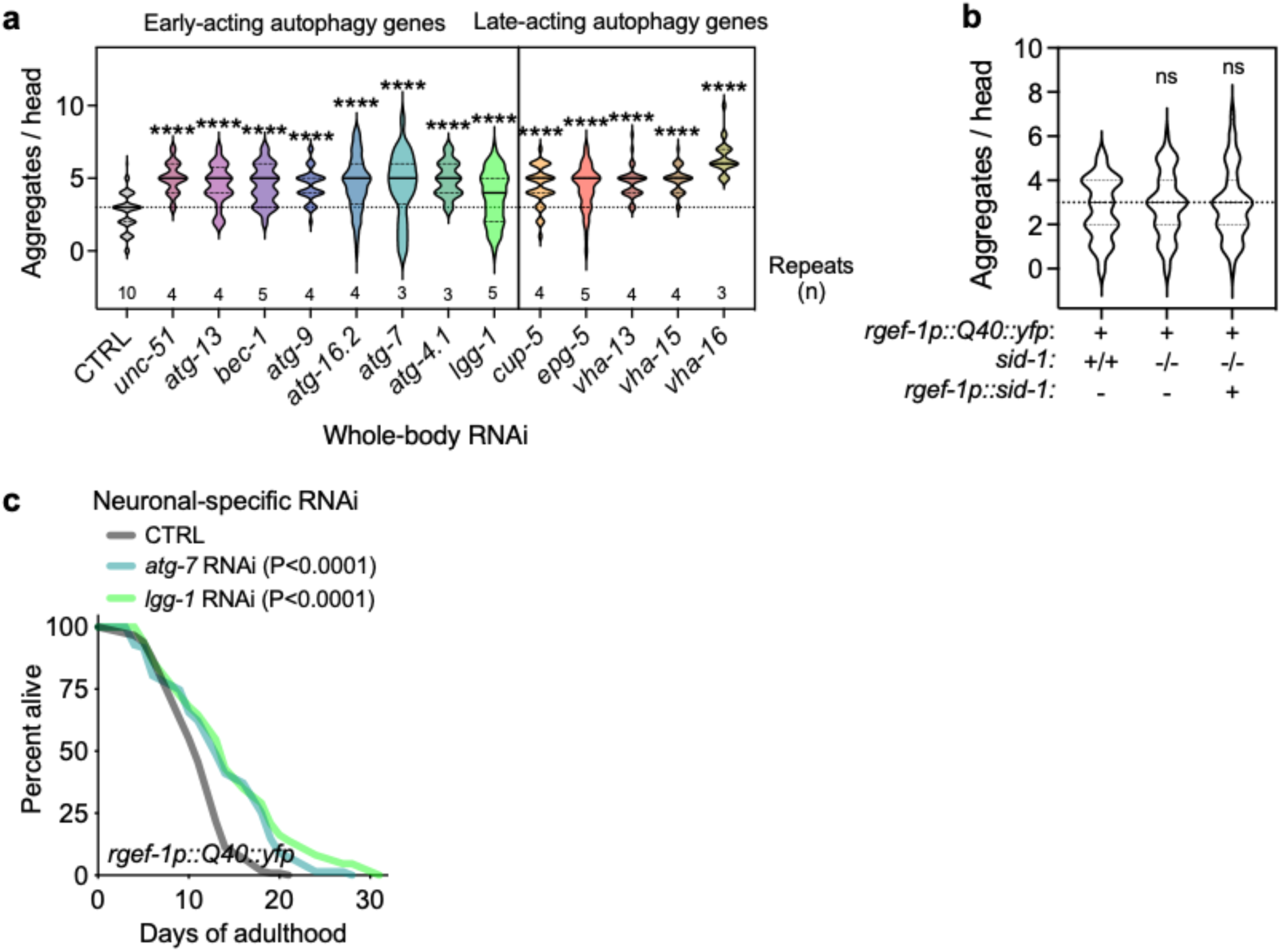
Neuronal PolyQ aggregation is increased by whole-body inhibition of autophagy genes, but unchanged in neuronal RNAi strains. (**A**) Neuronal PolyQ aggregates were scored in day 5 animals expressing *rgef-1::Q40::yfp* fed bacteria expressing dsRNAi for early- and late-acting autophagy genes from hatching on. Violin plots of data from indicated number of replicates, each with at least N=10 animals with solid line indicating median and dashed lines indicating quartiles ****P < 0.0001, by one-way ANOVA. **(B)** Neuronal PolyQ aggregates in day 7 animals expressing *rgef-1::Q40::yfp* in WT and *sid-1* mutants with or without *rgef-1p::sid-1* transgene. Violin plots of data from n=3 experiments, with N=41-48 animals with solid lines indicating median and dashed quartiles. ns P > 0.05 by one-way ANOVA. **(C)** Lifespan analysis of animals capable of neuronal-only RNAi and expressing neuronal PolyQ proteins (*sid-1; rgef-1p::sid-1 + rgef-1p::gfp; rgef-1::Q40::yfp*), which were fed bacteria expressing empty vector (CTRL), *atg-7,* or *lgg-1/Atg8* dsRNA from hatching on. Statistical significance determined by log-rank test. See **Table S3** for details and repeats.

**Figure S3.**
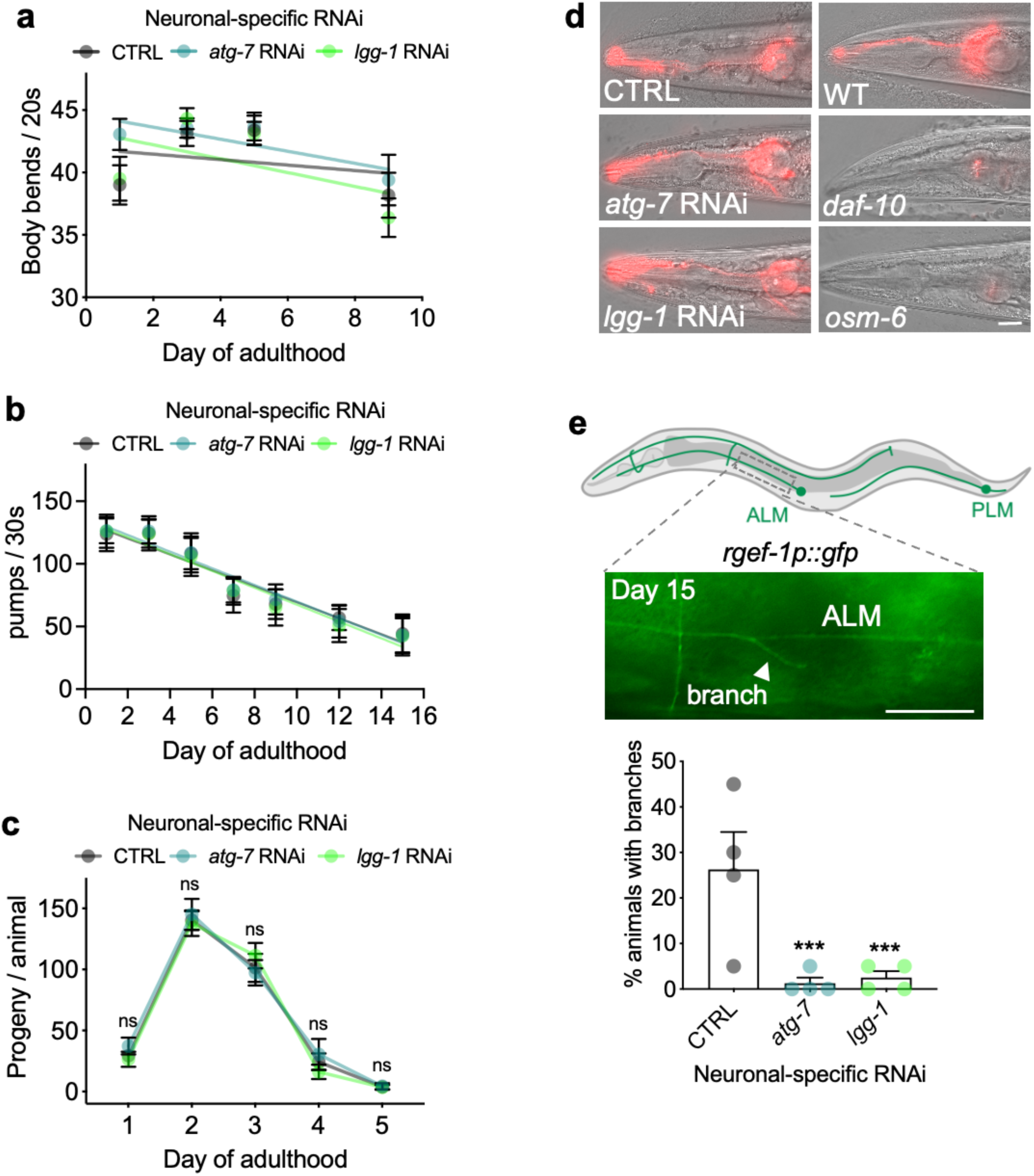
Healthspan and neuronal phenotypes of animals subjected to whole-life, neuronal inhibition of *atg-7* and *lgg-1/Atg8*. **(a)** Number of body bends per 20 s of animals capable of neuronal-only RNAi (*sid-1; rgef-1p::sid-1 + rgef-1p::gfp*) fed bacteria expressing empty vector (CTRL), *atg-7,* or *lgg-1/Atg8* dsRNA from hatching in liquid at different ages. Error bars indicate s.e.m. of one representative experiment, each with N=16 animals. Experiment was performed three times with similar results. Linear regression comparison versus CTRL: *atg-7* RNAi: P_slope_=0.5; P_y-intercept_=0.02; *lgg-1/Atg8* RNAi: P_slope_=0.3; P_y-intercept_=0.01. **(b)** For pharyngeal pumping assays, the number of contractions in the terminal pharyngeal bulb of animals capable of neuronal-specific RNAi (*sid-1; rgef-1p::sid-1 + rgef-1p::gfp*) fed bacteria expressing empty vector (CTRL), *atg-7,* or *lgg-1/Atg8* dsRNA from hatching was counted for 30 s. Error bars indicate s.d. of n=3 experiments, with N=30 animals. Linear regression comparison versus CTRL: *atg-7* RNAi: P_slope_=0.4; P_y-intercept_=0.4; *lgg-1* RNAi: P_slope_=0.3; P_y-intercept_=0.5. **(c)** For reproductive span, number of eggs and larvae produced per day per animal capable of neuronal-specific RNAi (*sid-1; rgef-1p::sid-1 + rgef-1p::gfp*) fed bacteria expressing empty vector (CTRL), *atg-7,* or *lgg-1/Atg8* dsRNA from hatching were counted. Error bars indicate s.d. of n=2 experiments, with N=18-22 animals. ns, P > 0.05, by two-way ANOVA. **(d)** Analysis of integrity of sensory neurons in animals capable of neuronal-only RNAi (*sid-1; rgef-1p::sid-1 + rgef-1p::gfp*) and fed bacteria expressing empty vector (CTRL), *atg-7,* or *lgg-1/Atg8* dsRNA from hatching on. Day 5 animals were soaked into DiI florescence red solution and analyzed for the ability to intake DiI to stain amphid and phasmid sensory neurons (see Methods). Sensory mutants *daf-10(e1387)* and *osm-6(p811)* were included in analysis as negative controls. Shown are representative images of n=3 experiments with N=10 animals each. Scale bar, 20 µm. (**d**) Neuronal branches originating from ALM and PLM neurons in animals capable of neuronal-only RNAi (*sid-1; rgef-1p::sid-1*) expressing *rgef-1p::gfp.* Representative image showing a branch (arrowhead) grew from ALM neuron on day 15. Scale bar: 20 µm. Quantification of percent of day 15 animals with branches fed bacteria expressing empty vector (CTRL), *atg-7*, or *lgg-1/Atg8* dsRNA from hatching. Error bars indicate s.d. of n=4 experiments, each with N=20 animals. ***P < 0.001, by Cochran-Mandel Hansel test.

**Figure S4.**
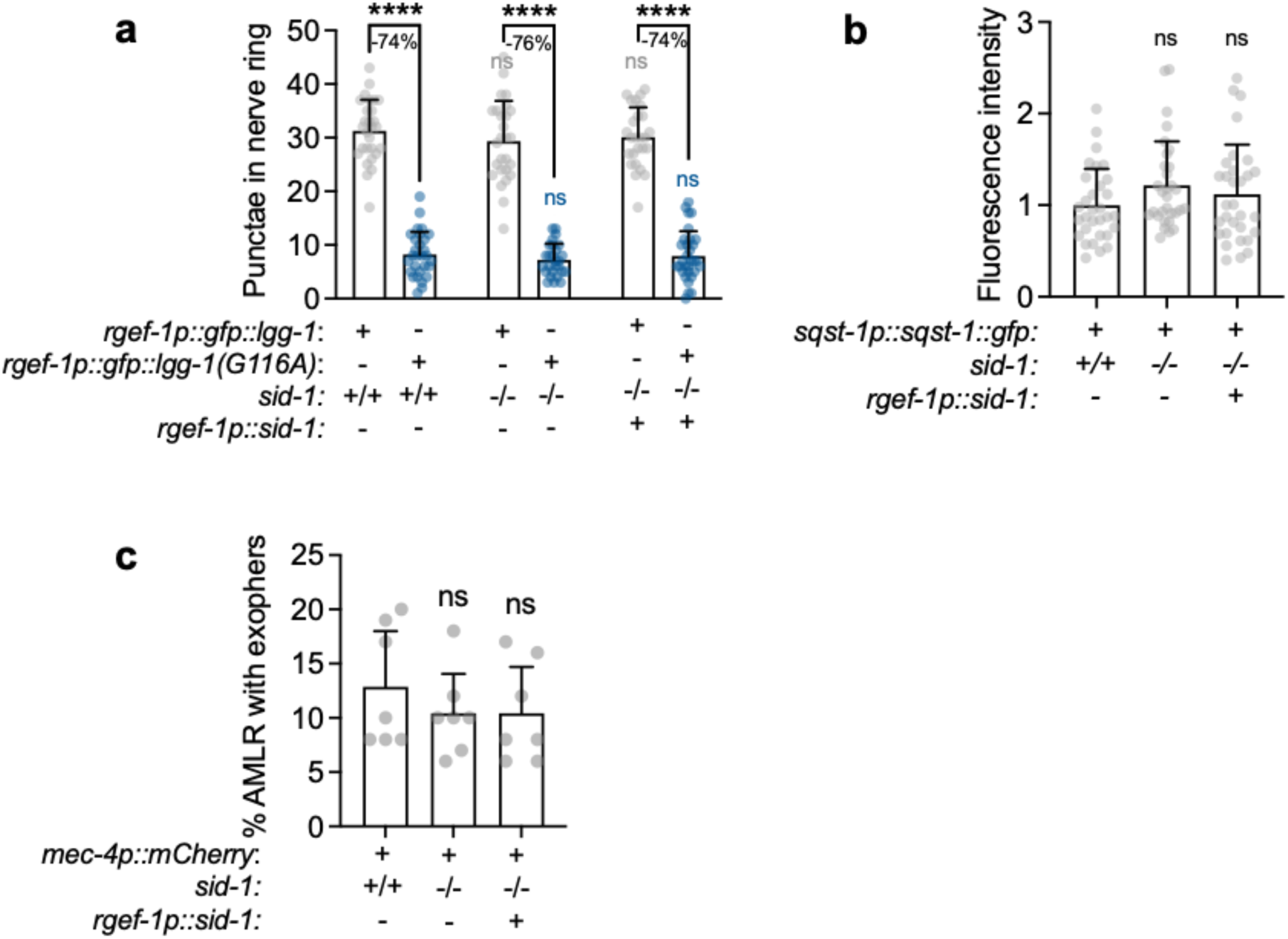
Neuronal autophagy status and exopher production are unchanged in animals expressing *sid-1* in neurons. **(a)** Neuronal GFP::LGG-1 and GFP::LGG-1(G116A) punctae were scored in day 1 wild-type (*sid-1*+/+) and *sid-1(qt9)* (*sid-1-/-)* animals with or without *rgef-1p::sid-1* transgene (*rgef-1p::sid-1* (+)). Error bars indicate s.d. of n=3 replicates with N=26-31 animals. Comparison between strains: ns P > 0.05, comparison of lipidated and unlipidated structures: ****P < 0.0001, by one-way ANOVA. **(B)** Fluorescence intensity of head region of animals expressing *sqst-1p::sqst-1::gfp* in wild-type (*sid-1*+/+), and *sid-1(qt9)* (*sid-1(-/-)*) animals with or without *rgef-1p::sid-1* transgene (*rgef-1p::sid-1* (+)) on day 1 of adulthood. Error bars indicate s.d. of n=3 with N=31 animals. ns P > 0.05 by one-way ANOVA. **(C)** Percent exophers in day 2 animals expressing *mec-4p::mCherry* in WT, and *sid-1(qt9)* mutants (*sid-1-/-)* with or without *rgef-1p::sid-1* transgene. Error bars indicate s.d. of n=7 experiments, each with N=25-62 animals. ns P > 0.05, by Cochran-Mandel Hansel test.

**Figure S5.**
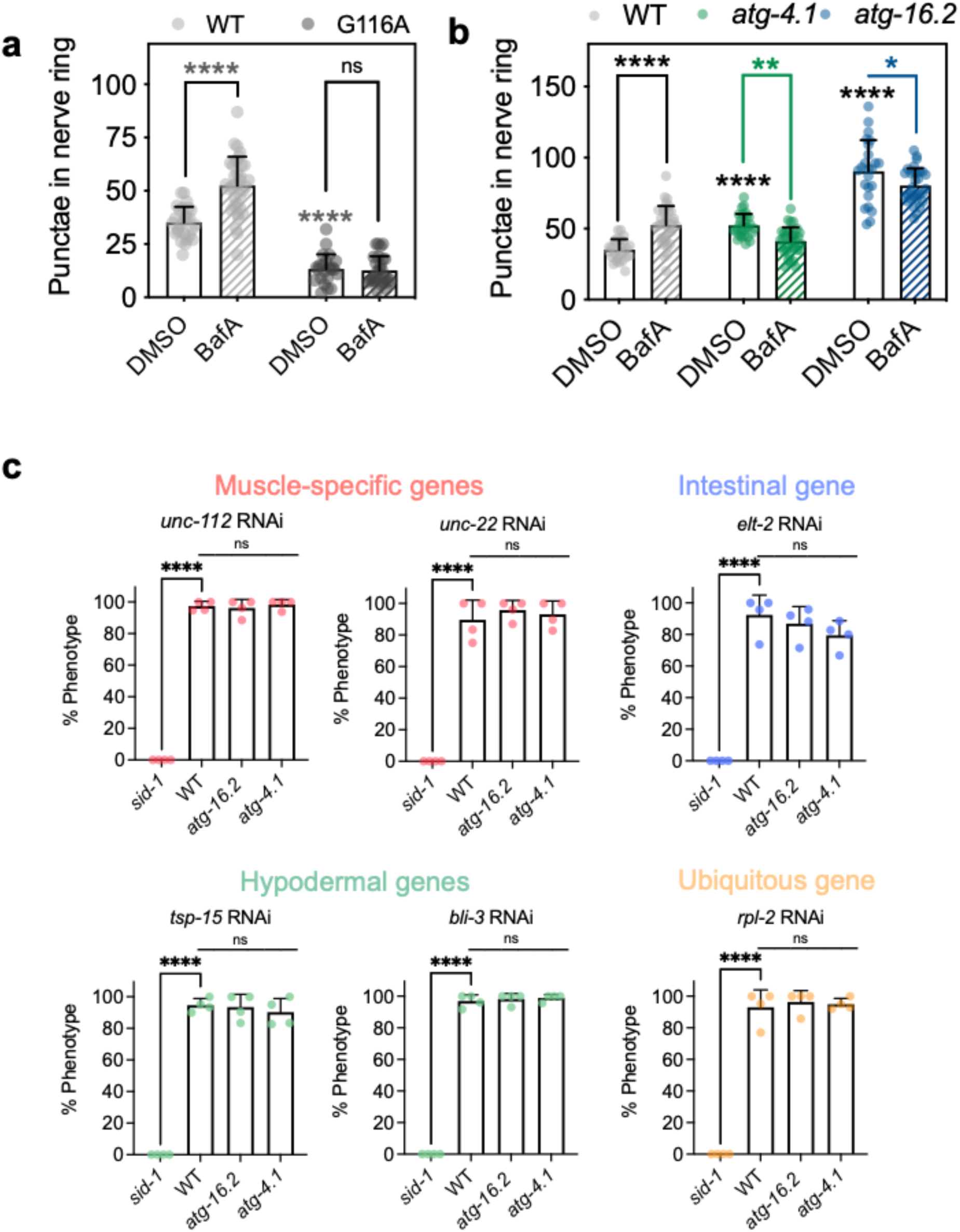
*atg-16.2* and *atg-4.1* mutants display similar autophagy- and RNAi phenotypes. (**A**) Autophagy flux in day 1 wild-type (WT) animals expressing either *rgef-tp::gfp::lgg-1* or *rgef-p::lgg-1(G116A).* GFP::LGG-1-positive punctae were quantified in nerve-ring neurons after injection of animals with vehicle (DMSO) or bafilomycin A1 (BafA) in n=3 with N=23-35 animals. Error bars indicate s.d. ns *P* > 0.05, \*\*\*\**P* < 0.0001, by two-way ANOVA with Tukey’s multiple comparisons test. (**B**) Autophagy flux in WT, *atg-4.1(bp501)*, and *atg-16.2(ok3224)* animals on day 1 of adulthood expressing *rgef-1p::gfp::lgg-1.* Day 1 animals were injected with vehicle (DMSO) or bafilomycin A1 (BafA) to block autophagy at the lysosomal acidification step and analyzed. GFP::LGG-1/Atg8-positive punctae were quantified in nerve-ring neurons in n=3 experiments with N=24-35 animals. Error bars indicate s.d. *P < 0.05, **P < 0.01, ****P < 0.0001, by two-way ANOVA with Tukey’s multiple comparisons test. (**C**) Phenotypes of day 2 *sid-1(qt9)*, WT, *atg-4.1(bp501)*, and *atg-16.2(ok3224)* animals fed bacteria expressing dsRNA against a specific gene expressing in different tissues from hatching on. Phenotypes for knockdown of each gene: Muscle-specific function: *unc-112*: paralysis, *unc-22*: twitching; Hypodermal-specific function: *tsp-15, bli-3*: blisters; intestinal-specific function: *elt-2*: small and clear, ubiquitous function: *rpl-2*: larval arrest. Error bars are s.d. of n=4 experiments with N=21-32 animals each. ns *P* > 0.05, \*\*\*\**P* < 0.0001, by one-way ANOVA.

**Figure S6.**
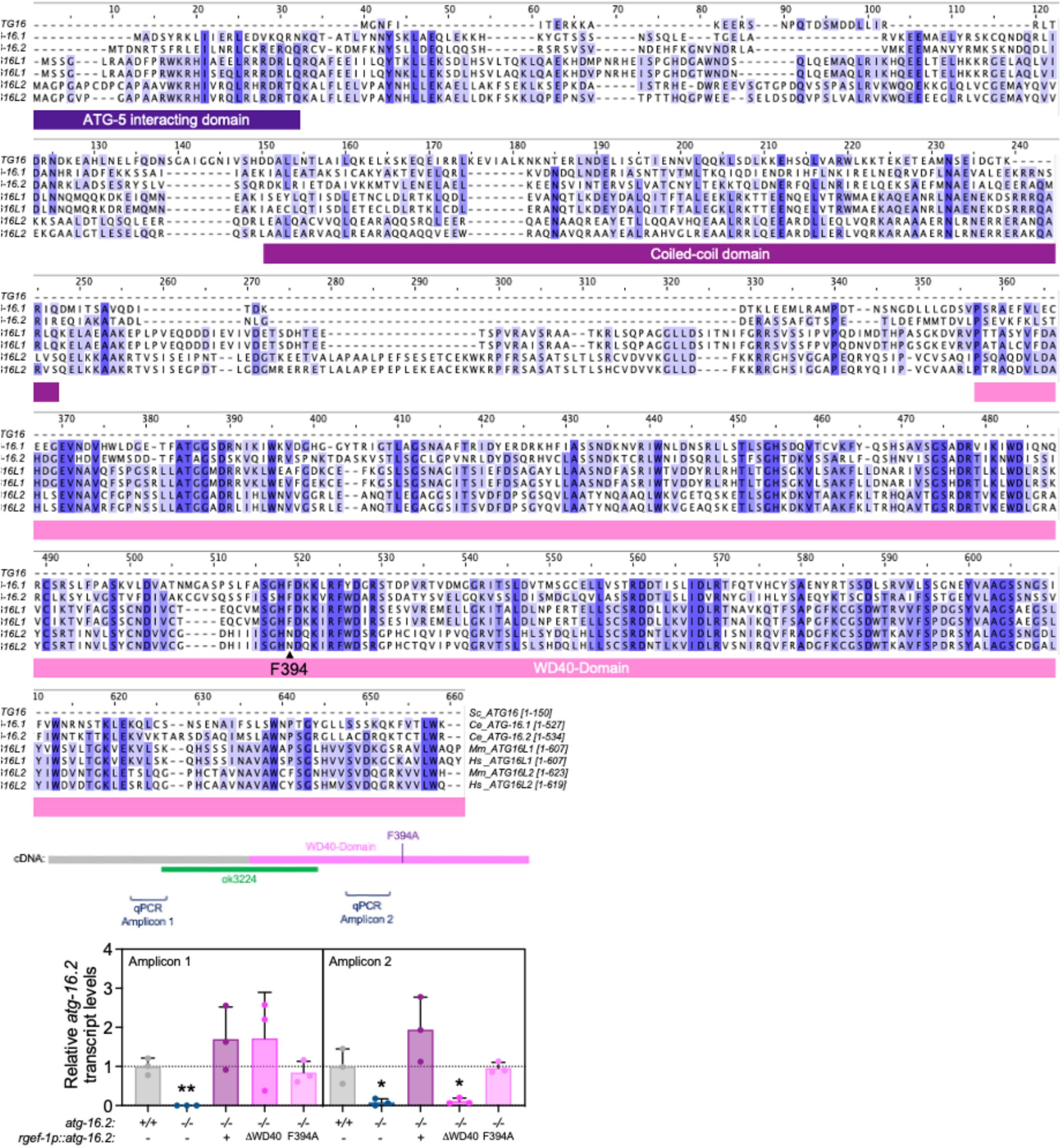
Multiple sequence alignment of ATG16 proteins and *atg-16.2* expression levels *legans*. (**A**) Primary sequences of ATG16 proteins from *S. cerevisiae* (1 isoform, Sc_ATG16), *C. elegans* (2 isoforms, Ce_ATG-16.1, Ce_ATG-16.2), *Mus musculus* (2 isoforms, Mm_ATG16L1, MmATG16L2), and humans (2 isoforms, Hs_ATG16L1, Hs_ATG16L2) were aligned by the Clustal Omega and colored by conservation with darker shades indicated increased conservation (Jalview). Protein elements of ATG16 proteins are drawn under the alignment and phenylalanine 349 is indicated with an arrow. Sequences showed are ATG16 (NP_013882.1) in *S. cerevisiae*, ATG16.2 (NP_495299.2) in *C. elegans*, ATG16 (NP_001138124.2) in *D. melanogaster*, ATG16L1 (NP_001192320.1) in *M. musculus*, and ATG16L1 (NP_001350671.1) in *H. sapiens*. (**B**) Transcript levels of *atg-16.2* in wild-type (WT), *atg-16.2(ok3224)*, and *atg-16.2(ok3224)* animals expressing full-length *atg-16.2*, *atg-16.2(ΔWD40)*, or *atg-16.2(F394A)* from the neuronal *rgef-1* promoter. Schematic of *atg-16.2* cDNA indicates the *ok3224* deletion, the WD40 domain, the position of the F394A point mutation, and the amplicons produced by the primers used in this experiment. Data are the mean and s.e.m. of three biological replicates, each with three technical replicates, and are normalized to the mean expression levels of three housekeeping genes. ns *P* > 0.05, \**P* < 0.05, \*\**P* < 0.01 by one-way ANOVA.

## Notes

### Competing Interest Statement

The authors have declared no competing interest.

