## Supplemental Tables for "Inhibition of early-acting autophagy genes in *C. elegans* neurons improves protein homeostasis, promotes exopher production, and extends lifespan via the ATG-16.2 WD40 domain"

**Supplementary Tables**

**Table S1: Summary of lifespan experiments of comparing wild-type, *sid-1,* and neuronal-specific RNAi strains**

| Exp # | Strain Name | Strain Genotype | MLS (Days) | % MLS change | P value | N animals/ total |
|---|---|---|---|---|---|---|
| 1 | N2 | wild type (WT) | 14.4 | | | 93/120 |
| | MAH346 | *sid-1* | 14.7 | 2 | 0.3 | 87/120 |
| | MAH676 | *sid-1; sqIs69[rgef-1p::sid-1+rgef-1p::gfp]* | 17.2 | 19 | 0.02 | 91/121 |
| | MAH677 | *sid-1; sqIs71[rgef-1p::sid-1+rgef-1p::gfp]* | 13.5 | -6 | 0.1 | 85/122 |
| 2 | N2 | WT | 13.0 | | | 96/120 |
| | MAH346 | *sid-1* | 13.3 | 2 | 0.4 | 100/120 |
| | MAH676 | *sid-1; sqIs69[rgef-1p::sid-1+rgef-1p::gfp]* | 15.0 | 15 | 0.03 | 113/120 |
| | MAH677 | *sid-1; sqIs71[rgef-1p::sid-1+rgef-1p::gfp]* | 14.8 | 14 | 0.02 | 74/120 |
| 3 | N2 | WT | 18.5 | | | 69/98 |
| | N2 | WT | 16.6 | -11 | 0.2 | 67/120 |
| | MAH346 | *sid-1* | 17.7 | -4 | 0.6 | 88/120 |
| | MAH676 | *sid-1; sqIs69[rgef-1p::sid-1+rgef-1p::gfp]* | 18.4 | -1 | 1.0 | 84/118 |
| | MAH677 | *sid-1; sqIs71[rgef-1p::sid-1+rgef-1p::gfp]* | 18.7 | 1 | 0.9 | 77/120 |
| | MAH656 | *sqIs69[rgef-1p::sid-1+rgef-1p::gfp]* | 18.6 | 1 | 1.0 | 92/120 |
| | MAH657 | *sqIs71[rgef-1p::sid-1+rgef-1p::gfp]* | 19.6 | 6 | 0.4 | 81/120 |
| | MAH630 | *sqEx101[rgef-1p::sid-1+rgef-1p::gfp]* | 16.1 | -13 | 0.02 | 83/120 |
| 4 | N2 | WT | 16.3 | | | 85/159 |
| | MAH677 | *sid-1; sqIs69[rgef-1p::sid-1+rgef-1p::gfp]* | 16.8 | 3 | 0.8 | 68/161 |
| 5 | N2 | WT | 19.1 | | | 50/120 |
| | N2 | WT | 19.8 | 4 | 0.6 | 47/115 |
| | MAH346 | *sid-1* | 18.5 | -3 | 0.5 | 43/121 |
| | MAH676 | *sid-1; sqIs69[rgef-1p::sid-1+rgef-1p::gfp]* | 20.6 | 8 | 0.5 | 56/116 |
| | MAH677 | *sid-1; sqIs71[rgef-1p::sid-1+rgef-1p::gfp]* | 20.8 | 9 | 0.4 | 49/120 |
| | MAH656 | *sqIs69[rgef-1p::sid-1+rgef-1p::gfp]* | 19.5 | 2 | 0.8 | 57/111 |
| | MAH657 | *sqIs71[rgef-1p::sid-1+rgef-1p::gfp]* | 19.6 | 3 | 0.9 | 53/121 |

| | | | | | | |
|---|---|---|---|---|---|---|
| | MAH630 | *sqEx101[rgef-1p::sid-1+rgef-1p::gfp]* | 17.4 | -9 | 0.1 | 60/115 |
| 6 | N2 | WT | 15.2 | | | 83/108 |
| | N2 | WT | 14.3 | -6 | 0.5 | 81/105 |
| | MAH346 | *sid-1* | 15.4 | 1 | 0.8 | 74/107 |
| | MAH676 | *sid-1; sqIs69[rgef-1p::sid-1+rgef-1p::gfp]* | 18.5 | 22 | 0.0001 | 80/108 |
| | MAH677 | *sid-1; sqIs71[rgef-1p::sid-1+rgef-1p::gfp]* | 15.6 | 3 | 0.5 | 73/108 |
| | MAH656 | *sqIs69[rgef-1p::sid-1+rgef-1p::gfp]* | 18.2 | 20 | <0.0001 | 90/111 |
| | MAH657 | *sqIs71[rgef-1p::sid-1+rgef-1p::gfp]* | 19.5 | 29 | <0.0001 | 94/112 |
| | MAH630 | *sqEx101[rgef-1p::sid-1+rgef-1p::gfp]* | 15.3 | 1 | 0.6 | 74/109 |
| 7 | N2 | WT | 15.1 | | | 78/120 |
| | MAH346 | *sid-1* | 14.4 | -5 | 0.7 | 61/121 |
| 8 | N2 | WT | 17.9 | | | 55/118 |
| | MAH346 | *sid-1* | 18.1 | 1.1 | 0.8 | 50/120 |
| 9 | N2 | *WT* | 16.7 | | | 89/119 |
| | MAH346 | *sid-1* | 18.3 | 10 | 0.003 | 109/122 |
| 10 | N2 | WT | 17.2 | | | 77/137 |
| | MAH677 | *sid-1; sqIs69[rgef-1p::sid-1+rgef-1p::gfp]* | 17.3 | 1 | 1.0 | 56/163 |
| 11 | N2 | WT | 14.3 | | | 114/120 |
| | MAH656 | *sqIs69[rgef-1p::sid-1+rgef-1p::gfp]* | 14.8 | 6 | 0.4 | 111/120 |
| | MAH677 | *sid-1; sqIs71[rgef-1p::sid-1+rgef-1p::gfp]* | 16.8 | 18 | 0.01 | 103/122 |
| 12 | N2 | WT | 17.5 | | | 108/114 |
| | MAH346 | *sid-1* | 20.1 | 15 | 0.001 | 109/113 |
| | MAH677 | *sid-1; sqIs71[rgef-1p::sid-1+rgef-1p::gfp]* | 16.4 | -6 | 0.4 | 89/107 |

**Supplementary Table 1:** Lifespan analysis of wild-type (WT) animals, *sid-1* mutants, WT animals expressing *rgef-1p::sid-1*, and *sid-1* mutants expressing *rgef-1p::sid-1* (Strain name and genotype indicated). Exp: experiment number; experiments not conducted in conjunction with experiments listed in other tables. MLS: mean lifespan; % MLS: percentage change in lifespan compared with WT control; P-values calculated by log-rank test. N: observed deaths/total number of animals.

**Table S2: Summary of lifespan experiments in wild type animals after neuronal knockdown of *lgg-1* and *atg-7***

| Condition | Exp # | Strain Name | Strain Genotype | RNAi treat-ment | RNAi MLS (Days) | Control MLS (Days) | % MLS change | P value | N animals/ total |
|---|---|---|---|---|---|---|---|---|---|
| *lgg-1* RNAi (adult-only) | 1 | MAH677 | *sid-1; sqIs71[rgef-1p::sid-1+rgef-1p::gfp]* | *lgg-1* | 20.1 | 17.8 | 37.0 | <0.0001 | 98/108 |
| | 2 | MAH676 | *sid-1; sqIs69[rgef-1p::sid-1+rgef-1p::gfp]* | *lgg-1* | 21.9 | 17.0 | 29.0 | <0.0001 | 90/120 |
| | 3 | MAH676 | *sid-1; sqIs69[rgef-1p::sid-1+rgef-1p::gfp]* | *lgg-1* | 25.7 | 17.8 | 44.0 | <0.0001 | 89/103 |
| | 4 | MAH677 | *sid-1; sqIs71[rgef-1p::sid-1+rgef-1p::gfp]* | *lgg-1* | 18.1 | 16.1 | 12.4 | 0.03 | 54/105 |
| | 5 | MAH677 | *sid-1; sqIs71[rgef-1p::sid-1+rgef-1p::gfp]* | *lgg-1* | 23.9 | 18.6 | 28.5 | <0.0001 | 89/126 |
| | 6 | MAH677 | *sid-1; sqIs71[rgef-1p::sid-1+rgef-1p::gfp]* | *lgg-1* | 21.1 | 16.1 | 31.1 | <0.0001 | 71/120 |
| | 7 | MAH677 | *sid-1; sqIs71[rgef-1p::sid-1+rgef-1p::gfp]* | *lgg-1* | 24.7 | 20.6 | 19.9 | <0.0001 | 79/120 |
| | 8 | MAH677 | *sid-1; sqIs71[rgef-1p::sid-1+rgef-1p::gfp]* | *lgg-1* | 23.1 | 18.8 | 22.3 | <0.0001 | 75/120 |
| | 9 | MAH677 | *sid-1; sqIs71[rgef-1p::sid-1+rgef-1p::gfp]* | *lgg-1* | 21.1 | 17.5 | 20.6 | 0.0002 | 93/114 |
| | 10 | MAH677 | *sid-1; sqIs71[rgef-1p::sid-1+rgef-1p::gfp]* | *lgg-1* | 22.3 | 19.4 | 14.9 | 0.001 | 80/101 |
| | 11 | MAH677 | *sid-1; sqIs71[rgef-1p::sid-1+rgef-1p::gfp]* | *lgg-1* | 22.9 | 16.8 | 36.3 | <0.0001 | 94/120 |
| | 12 | MAH677 | *sid-1; sqIs71[rgef-1p::sid-1+rgef-1p::gfp]* | *lgg-1* | 22.8 | 18.3 | 24.5 | <0.0001 | 96/122 |
| | 13 | MAH677 | *sid-1; sqIs71[rgef-1p::sid-1+rgef-1p::gfp]* | *lgg-1* | 21.9 | 20.0 | 9.5 | 0.06 | 82/109 |
| | 14 | MAH677 | *sid-1; sqIs71[rgef-1p::sid-1+rgef-1p::gfp]* | *lgg-1* | 21.2 | 18.7 | 13.4 | 0.01 | 71/106 |
| | 15 | MAH677 | *sid-1; sqIs71[rgef-1p::sid-1+rgef-1p::gfp]* | *lgg-1* | 22.7 | 16.9 | 34.0 | <0.0001 | 80/123 |
| | 16 | MAH677 | *sid-1; sqIs71[rgef-1p::sid-1+rgef-1p::gfp]* | *lgg-1* | 25.4 | 20.1 | 26.0 | <0.0001 | 89/120 |
| | 17 | MAH677 | *sid-1; sqIs71[rgef-1p::sid-1+rgef-1p::gfp]* | *lgg-1* | 23.4 | 21.2 | 10.4 | 0.0006 | 89/119 |
| | 18 | MAH677 | *sid-1; sqIs71[rgef-1p::sid-1+rgef-1p::gfp]* | *lgg-1* | 24.5 | 19.6 | 25.0 | <0.0001 | 81/108 |
| | 5 | MAH757 | *sid-1; sqEx101[rgef-1p::sid-1 + rgef-1p::gfp]* | *lgg-1* | 22.9 | 17.3 | 32.4 | <0.0001 | 93/118 |
| | 15 | MAH798 | *sid-1; sqEx131[rgef-1p::sid-1 + unc-122p::rfp]* | *lgg-1* | 20.4 | 17.4 | 17.2 | 0.0006 | 74/119 |
| | 16 | MAH848 | *sid-1; sqEx148[rgef-1p::sid-1 + rol-6]* | *lgg-1* | 23.4 | 19.1 | 22.5 | <0.0001 | 80/121 |
| | 19 | MAH677 | *sid-1; sqIs71[rgef-1p::sid-1+rgef-1p::gfp]* | *lgg-1* | 20.2 | 16.1 | 25.5 | <0.0001 | 75/120 |
| | 20 | MAH677 | *sid-1; sqIs71[rgef-1p::sid-1+rgef-1p::gfp]* | *lgg-1* | 22.3 | 19.0 | 17.4 | <0.0001 | 71/105 |
| | 21 | MAH677 | *sid-1; sqIs71[rgef-1p::sid-1+rgef-1p::gfp]* | *lgg-1* | 24.7 | 21.4 | 15.4 | 0.004 | 61/105 |
| *lgg-1* RNAi (whole-life) | 2 | MAH676 | *sid-1; sqIs69[rgef-1p::sid-1+rgef-1p::gfp]* | *lgg-1* | 24.1 | 17.0 | 42.0 | <0.0001 | 107/120 |
| | 22 | MAH677 | *sid-1; sqIs71[rgef-1p::sid-1+rgef-1p::gfp]* | *lgg-1* | 22.9 | 19.9 | 15.1 | <0.0001 | 95/120 |
| | 23 | MAH677 | *sid-1; sqIs71[rgef-1p::sid-1+rgef-1p::gfp]* | *lgg-1* | 22.2 | 19.8 | 12.1 | 0.02 | 82/101 |
| | 24 | MAH677 | *sid-1; sqIs71[rgef-1p::sid-1+rgef-1p::gfp]* | *lgg-1* | 24.6 | 19.3 | 27.5 | <0.0001 | 101/117 |

| | | | | | | | | | |
|---|---|---|---|---|---|---|---|---|---|
| | 25 | MAH677 | *sid-1; sqIs71[rgef-1p::sid-1+rgef-1p::gfp]* | *lgg-1* | 26.3 | 21.2 | 24.1 | <0.0001 | 78/101 |
| | 26 | MAH677 | *sid-1; sqIs71[rgef-1p::sid-1+rgef-1p::gfp]* | *lgg-1* | 25.6 | 22.1 | 29.7 | <0.0001 | 79/120 |
| | 27 | MAH677 | *sid-1; sqIs71[rgef-1p::sid-1+rgef-1p::gfp]* | *lgg-1* | 27.3 | 20.9 | 30.6 | <0.0001 | 72/120 |
| | 28 | MAH677 | *sid-1; sqIs71[rgef-1p::sid-1+rgef-1p::gfp]* | *lgg-1* | 25.1 | 19.5 | 28.7 | <0.0001 | 87/124 |
| | 29 | MAH677 | *sid-1; sqIs71[rgef-1p::sid-1+rgef-1p::gfp]* | *lgg-1* | 26.3 | 20.2 | 30.1 | <0.0001 | 61/119 |
| | 30 | MAH798 | *sid-1; sqEx131[rgef-1p::sid-1 + unc-122p::rfp]* | *lgg-1* | 17.8 | 15.4 | 15.6 | 0.001 | 74/119 |
| | 31 | MAH677 | *sid-1; sqIs71[rgef-1p::sid-1+rgef-1p::gfp]* | *lgg-1* | 26.4 | 18.1 | 44.2 | <0.0001 | 80/120 |
| | 32 | MAH798 | *sid-1; sqEx131[rgef-1p::sid-1 + unc-122p::rfp]* | *lgg-1* | 21.8 | 17.9 | 21.8 | <0.0001 | 67/108 |
| | 33 | MAH798 | *sid-1; sqEx131[rgef-1p::sid-1 + unc-122p::rfp]* | *lgg-1* | 25.7 | 16.4 | 56.7 | <0.0001 | 62/111 |
| | 34 | MAH798 | *sid-1; sqEx131[rgef-1p::sid-1 + unc-122p::rfp]* | *lgg-1* | 21.7 | 16.1 | 34.7 | <0.0001 | 91/120 |
| | 35 | MAH798 | *sid-1; sqEx131[rgef-1p::sid-1 + unc-122p::rfp]* | *lgg-1* | 19.3 | 17.8 | 8.4 | 0.02 | 91/120 |
| | 36 | MAH677 | *sid-1; sqIs71[rgef-1p::sid-1+rgef-1p::gfp]* | *lgg-1* | 27.7 | 24.3 | 14.0 | <0.0001 | 89/96 |
| | 37* | MAH677 | *sid-1; sqIs71[rgef-1p::sid-1+rgef-1p::gfp]* | *lgg-1* | 21.5 | 26.2 | 21.9 | <0.0001 | 99/108 |
| | 38 | MAH677 | *sid-1; sqIs71[rgef-1p::sid-1+rgef-1p::gfp]* | *lgg-1* | 23.8 | 25.6 | 7.0 | 0.003 | 107/133 |
| *lgg-1* RNAi (L1) | 39 | MAH676 | *sid-1; sqIs69[rgef-1p::sid-1+rgef-1p::gfp]* | *lgg-1* | 23.4 | 18.5 | 26.0 | <0.0001 | 116/120 |
| | 40 | MAH677 | *sid-1; sqIs71[rgef-1p::sid-1+rgef-1p::gfp]* | *lgg-1* | 22.7 | 18.1 | 26.0 | <0.0001 | 105/120 |
| | 56 | MAH677 | *sid-1; sqIs71[rgef-1p::sid-1+rgef-1p::gfp]* | *lgg-1* | 23.5 | 20.6 | 14.0 | 0.01 | 97/102 |
| | 1 | MAH677 | *sid-1; sqIs71[rgef-1p::sid-1+rgef-1p::gfp]* | *lgg-1* | 23.7 | 17.8 | 33.0 | <0.0001 | 75/91 |
| | 41 | MAH676 | *sid-1; sqIs69[rgef-1p::sid-1+rgef-1p::gfp]* | *lgg-1* | 21.6 | 16.8 | 29.0 | <0.0001 | 87/104 |
| | 3 | MAH676 | *sid-1; sqIs69[rgef-1p::sid-1+rgef-1p::gfp]* | *lgg-1* | 23.5 | 17.8 | 32.0 | <0.0001 | 68/101 |
| *atg-7* RNAi (adult-only) | 6 | MAH677 | *sid-1; sqIs71[rgef-1p::sid-1+rgef-1p::gfp]* | *atg-7* | 17.6 | 16.1 | 9.3 | 0.06 | 59/128 |
| | 7 | MAH677 | *sid-1; sqIs71[rgef-1p::sid-1+rgef-1p::gfp]* | *atg-7* | 23.2 | 20.6 | 12.6 | 0.01 | 72/119 |
| | 17 | MAH677 | *sid-1; sqIs71[rgef-1p::sid-1+rgef-1p::gfp]* | *atg-7* | 22.8 | 21.2 | 7.5 | 0.005 | 89/117 |
| | 19 | MAH677 | *sid-1; sqIs71[rgef-1p::sid-1+rgef-1p::gfp]* | *atg-7* | 21.4 | 16.1 | 32.9 | <0.0001 | 75/120 |
| | 18 | MAH677 | *sid-1; sqIs71[rgef-1p::sid-1+rgef-1p::gfp]* | *atg-7* | 22.6 | 19.6 | 15.3 | <0.0001 | 97/108 |
| | 4 | MAH677 | *sid-1; sqIs71[rgef-1p::sid-1+rgef-1p::gfp]* | *atg-7* | 19.1 | 16.1 | 18.6 | 0.001 | 65/99 |
| | 21 | MAH677 | *sid-1; sqIs71[rgef-1p::sid-1+rgef-1p::gfp]* | *atg-7* | 26.4 | 21.4 | 23.4 | 0.0001 | 46/73 |
| *atg-7* RNAi (whole-life) | 23 | MAH677 | *sid-1; sqIs71[rgef-1p::sid-1+rgef-1p::gfp]* | *atg-7* | 22.2 | 19.8 | 12.1 | 0.001 | 104/118 |
| | 24 | MAH677 | *sid-1; sqIs71[rgef-1p::sid-1+rgef-1p::gfp]* | *atg-7* | 21.6 | 19.3 | 11.9 | 0.001 | 102/120 |
| | 25 | MAH677 | *sid-1; sqIs71[rgef-1p::sid-1+rgef-1p::gfp]* | *atg-7* | 25.1 | 21.2 | 18.4 | <0.0001 | 94/111 |
| | 26 | MAH677 | *sid-1; sqIs71[rgef-1p::sid-1+rgef-1p::gfp]* | *atg-7* | 25 | 22.1 | 34.8 | 0.0006 | 73/120 |

| | | | | | | | | | |
|---|---|---|---|---|---|---|---|---|---|
| | 27 | MAH677 | *sid-1; sqIs71[rgef-1p::sid-1+rgef-1p::gfp]* | *atg-7* | 24.5 | 20.9 | 17.2 | 0.0006 | 94/120 |
| | 29 | MAH677 | sid-1; sqIs71[rgef-1p::sid-1+rgef-1p::gfp] | atg-7 | 24.3 | 20.2 | 20.3 | <0.0001 | 79/113 |
| | 30 | MAH798 | *sid-1; sqEx131[rgef-1p::sid-1 + unc-122p::rfp]* | *atg-7* | 18.7 | 15.4 | 21.4 | <0.0001 | 89/120 |
| | 31 | MAH677 | *sid-1; sqIs71[rgef-1p::sid-1+rgef-1p::gfp]* | *atg-7* | 24.4 | 18.1 | 34.8 | <0.0001 | 87/120 |
| | 32 | MAH798 | *sid-1; sqEx131[rgef-1p::sid-1 + unc-122p::rfp]* | *atg-7* | 20.8 | 17.9 | 16.2 | 0.0001 | 74/112 |
| | 33 | MAH798 | *sid-1; sqEx131[rgef-1p::sid-1 + unc-122p::rfp]* | *atg-7* | 20.9 | 16.4 | 27.4 | <0.0001 | 54/118 |
| | 34 | MAH798 | *sid-1; sqEx131[rgef-1p::sid-1 + unc-122p::rfp]* | *atg-7* | 21.3 | 16.1 | 32.2 | <0.0001 | 96/120 |
| | 35 | MAH798 | *sid-1; sqEx131[rgef-1p::sid-1 + unc-122p::rfp]* | *atg-7* | 19.7 | 17.8 | 10.7 | 0.01 | 96/120 |
| | 38 | MAH677 | *sid-1; sqIs71[rgef-1p::sid-1+rgef-1p::gfp]* | *atg-7* | 23.8 | 24.7 | 4.0 | 0.2 | 117/140 |
| | 37* | MAH677 | *sid-1; sqIs71[rgef-1p::sid-1+rgef-1p::gfp]* | *atg-7* | 21.5 | 26.2 | 21.9 | <0.0001 | 92/108 |
| | 36 | MAH677 | *sid-1; sqIs71[rgef-1p::sid-1+rgef-1p::gfp]* | *atg-7* | 28.5 | 24.3 | 17.0 | <0.0001 | 88/96 |
| *atg-7* RNAi (L1) | 39 | MAH676 | *sid-1; sqIs69[rgef-1p::sid-1+rgef-1p::gfp]* | *atg-7* | 21.3 | 18.5 | 15.0 | 0.0001 | 115/121 |
| | 40 | MAH677 | *sid-1; sqIs71[rgef-1p::sid-1+rgef-1p::gfp]* | *atg-7* | 22.1 | 18.1 | 22.0 | <0.0001 | 109/120 |
| | 41 | MAH676 | *sid-1; sqIs69[rgef-1p::sid-1+rgef-1p::gfp]* | *atg-7* | 20.8 | 16.8 | 24.0 | <0.0001 | 88/101 |
| *bec-1* RNAi (adult-only) | 42 | MAH676 | *sid-1; sqIs69[rgef-1p::sid-1+rgef-1p::gfp]* | *bec-1* | 19.7 | 21.1 | -7.0 | 0.07 | 80/99 |
| | 4 | MAH677 | *sid-1; sqIs71[rgef-1p::sid-1+rgef-1p::gfp]* | *bec-1* | 21.3 | 16.1 | 32.2 | <0.0001 | 56/106 |
| | 6 | MAH677 | *sid-1; sqIs71[rgef-1p::sid-1+rgef-1p::gfp]* | *bec-1* | 17.9 | 16.1 | 11.2 | 0.02 | 65/118 |
| | 17 | MAH677 | *sid-1; sqIs71[rgef-1p::sid-1+rgef-1p::gfp]* | *bec-1* | 22.1 | 21.2 | 4.3 | 0.06 | 100/122 |
| | 18 | MAH677 | *sid-1; sqIs71[rgef-1p::sid-1+rgef-1p::gfp]* | *bec-1* | 24.1 | 19.6 | 23.0 | <0.0001 | 90/108 |
| *bec-1* RNAi (whole-life) | 22 | MAH677 | *sid-1; sqIs71[rgef-1p::sid-1+rgef-1p::gfp]* | *bec-1* | 21.1 | 19.9 | 6.1 | 0.08 | 64/121 |
| *bec-1* RNAi (L1) | 39 | MAH676 | *sid-1; sqIs69[rgef-1p::sid-1+rgef-1p::gfp]* | *bec-1* | 18.8 | 18.5 | 2.0 | 0.4 | 114/123 |
| | 40 | MAH677 | *sid-1; sqIs71[rgef-1p::sid-1+rgef-1p::gfp]* | *bec-1* | 22.3 | 18.1 | 23.0 | <0.0001 | 109/120 |
| | 41 | MAH676 | *sid-1; sqIs69[rgef-1p::sid-1+rgef-1p::gfp]* | *bec-1* | 20.9 | 16.8 | 24.0 | <0.0001 | 88/102 |
| *atg-9* RNAi (adult-only) | 43 | MAH677 | *sid-1; sqIs71[rgef-1p::sid-1+rgef-1p::gfp]* | *atg-9* | 25.9 | 23.3 | 11.2 | 0.01 | 106/117 |
| *atg-9* RNAi (L1) | 39 | MAH676 | *sid-1; sqIs69[rgef-1p::sid-1+rgef-1p::gfp]* | *atg-9* | 21.2 | 18.5 | 15.0 | 0.0005 | 111/120 |
| | 40 | MAH677 | *sid-1; sqIs71[rgef-1p::sid-1+rgef-1p::gfp]* | *atg-9* | 21.6 | 18.1 | 20.0 | <0.0001 | 111/120 |
| | 41 | MAH676 | *sid-1; sqIs69[rgef-1p::sid-1+rgef-1p::gfp]* | *atg-9* | 20.8 | 16.8 | 24.0 | <0.0001 | 92/103 |
| *atg-13* RNAi (adult-only) | 9 | MAH677 | *sid-1; sqIs71[rgef-1p::sid-1+rgef-1p::gfp]* | *atg-13* | 24.6 | 17.5 | 40.6 | <0.0001 | 94/118 |
| | 10 | MAH677 | *sid-1; sqIs71[rgef-1p::sid-1+rgef-1p::gfp]* | *atg-13* | 23.2 | 19.4 | 19.6 | <0.0001 | 90/106 |

| | | | | | | | | | |
|---|---|---|---|---|---|---|---|---|---|
| | 11 | MAH677 | *sid-1; sqIs71[rgef-1p::sid-1+rgef-1p::gfp]* | *atg-13* | 19.6 | 16.8 | 16.7 | 0.003 | 79/118 |
| | 12 | MAH677 | *sid-1; sqIs71[rgef-1p::sid-1+rgef-1p::gfp]* | *atg-13* | 23.2 | 18.3 | 26.7 | <0.0001 | 102/120 |
| | 13 | MAH677 | *sid-1; sqIs71[rgef-1p::sid-1+rgef-1p::gfp]* | *atg-13* | 22.9 | 20 | 14.5 | 0.005 | 92/109 |
| | 14 | MAH677 | *sid-1; sqIs71[rgef-1p::sid-1+rgef-1p::gfp]* | *atg-13* | 22.0 | 18.7 | 17.6 | 0.0004 | 78/104 |
| *unc-51* RNAi (adult-only) | 1 | MAH677 | *sid-1; sqIs71[rgef-1p::sid-1+rgef-1p::gfp]* | *unc-51* | 21.3 | 17.8 | 20.0 | 0.003 | 97/106 |
| | 9 | MAH677 | *sid-1; sqIs71[rgef-1p::sid-1+rgef-1p::gfp]* | *unc-51* | 20.7 | 17.5 | 18.3 | 0.0008 | 95/120 |
| | 10 | MAH677 | *sid-1; sqIs71[rgef-1p::sid-1+rgef-1p::gfp]* | *unc-51* | 19.8 | 19.4 | 2.1 | 0.4 | 76/106 |
| | 11 | MAH677 | *sid-1; sqIs71[rgef-1p::sid-1+rgef-1p::gfp]* | *unc-51* | 20.5 | 16.8 | 22.1 | 0.0002 | 75/120 |
| | 19 | MAH677 | *sid-1; sqIs71[rgef-1p::sid-1+rgef-1p::gfp]* | *unc-51* | 17.2 | 16.1 | 6.8 | 0.1 | 76/119 |
| | 13 | MAH677 | *sid-1; sqIs71[rgef-1p::sid-1+rgef-1p::gfp]* | *unc-51* | 23.2 | 20 | 16.0 | 0.003 | 90/106 |
| *unc-51* RNAi (L1) | 1 | MAH677 | *sid-1; sqIs71[rgef-1p::sid-1+rgef-1p::gfp]* | *unc-51* | 20.1 | 17.8 | 13.0 | 0.03 | 79/98 |
| *atg-4.1* RNAi (whole-life) | 31 | MAH798 | *sid-1; sqEx131[rgef-1p::sid-1 + unc-122p::rfp]* | *atg-4.1* | 22.4 | 18.1 | 23.7 | <0.0001 | 76/119 |
| | 34 | MAH798 | *sid-1; sqEx131[rgef-1p::sid-1 + unc-122p::rfp]* | *atg-4.1* | 18.8 | 16.1 | 16.7 | 0.0002 | 88/120 |
| | 35 | MAH798 | *sid-1; sqEx131[rgef-1p::sid-1 + unc-122p::rfp]* | *atg-4.1* | 19.2 | 17.8 | 7.9 | 0.04 | 88/120 |
| *atg-16.2* RNAi (adult-only) | 45 | MAH677 | *sid-1; sqIs71[rgef-1p::sid-1+rgef-1p::gfp]* | *atg-16.2* | 16.3 | 16.8 | -2.9 | 0.6 | 63/120 |
| | 46 | MAH677 | *sid-1; sqIs71[rgef-1p::sid-1+rgef-1p::gfp]* | *atg-16.2* | 21.4 | 23.3 | -9.1 | 0.1 | 113/113 |
| *atg-16.2* RNAi (whole-life) | 47 | MAH677 | *sid-1; sqIs71[rgef-1p::sid-1+rgef-1p::gfp]* | *atg-16.2* | 20.5 | 26.5 | -22.6 | <0.0001 | 93/105 |
| | 37* | MAH677 | *sid-1; sqIs71[rgef-1p::sid-1+rgef-1p::gfp]* | *atg-16.2* | 21.5 | 20.3 | -5.6 | 0.07 | 100/110 |
| | 38 | MAH677 | *sid-1; sqIs71[rgef-1p::sid-1+rgef-1p::gfp]* | *atg-16.2* | 23.8 | 23.1 | -3.0 | 0.2 | 117/141 |
| | 48 | MAH677 | *sid-1; sqIs71[rgef-1p::sid-1+rgef-1p::gfp]* | *atg-16.2* | 21.7 | 22.8 | -4.8 | 0.18 | 112/140 |
| epg-5 RNAi (adult-only) | 49 | MAH677 | *sid-1; sqIs71[rgef-1p::sid-1+rgef-1p::gfp]* | *epg-5* | 22.3 | 20 | 6.7 | 0.3 | 52/100 |
| | 6 | MAH677 | *sid-1; sqIs71[rgef-1p::sid-1+rgef-1p::gfp]* | *epg-5* | 16.9 | 16.1 | 4.9 | 0.07 | 61/121 |
| | 4 | MAH677 | *sid-1; sqIs71[rgef-1p::sid-1+rgef-1p::gfp]* | *epg-5* | 17.8 | 16.1 | 10.5 | 0.1 | 45/94 |
| *epg-5* RNAi (whole-life) | 22 | MAH677 | *sid-1; sqIs71[rgef-1p::sid-1+rgef-1p::gfp]* | *epg-5* | 20.6 | 19.9 | 3.5 | 0.2 | 96/120 |
| | 50 | MAH677 | *sid-1; sqIs71[rgef-1p::sid-1+rgef-1p::gfp]* | *epg-5* | 15.9 | 18.4 | -13.9 | 0.0008 | 105/120 |
| *cup-5* RNAi (adult-only) | 49 | MAH677 | *sid-1; sqIs71[rgef-1p::sid-1+rgef-1p::gfp]* | *cup-5* | 21.3 | 20 | 1.9 | 0.8 | 52/107 |
| | 6 | MAH677 | *sid-1; sqIs71[rgef-1p::sid-1+rgef-1p::gfp]* | *cup-5* | 16.9 | 16.1 | 4.9 | 0.3 | 55/118 |
| | 4 | MAH677 | *sid-1; sqIs71[rgef-1p::sid-1+rgef-1p::gfp]* | *cup-5* | 16.7 | 16.1 | 3.7 | 0.3 | 68/114 |

| | | | | | | | | | |
|---|---|---|---|---|---|---|---|---|---|
| *cup-5* RNAi (whole-life) | 22 | MAH677 | *sid-1; sqIs71[rgef-1p::sid-1+rgef-1p::gfp]* | *cup-5* | 20.1 | 19.9 | 1.1 | 0.9 | 89/122 |
| | 50 | MAH677 | *sid-1; sqIs71[rgef-1p::sid-1+rgef-1p::gfp]* | *cup-5* | 19.3 | 18.4 | 4.8 | 0.8 | 103/120 |
| *vha-13* RNAi (adult-only) | 45 | MAH677 | *sid-1; sqIs71[rgef-1p::sid-1+rgef-1p::gfp]* | *vha-13* | 16.7 | 16.8 | -0.6 | 0.6 | 57/120 |
| | 49 | MAH677 | *sid-1; sqIs71[rgef-1p::sid-1+rgef-1p::gfp]* | *vha-13* | 21.0 | 20 | 0.5 | 0.6 | 59/109 |
| *vha-13* RNAi (whole-life) | 50 | MAH677 | *sid-1; sqIs71[rgef-1p::sid-1+rgef-1p::gfp]* | *vha-13* | 16.8 | 18.4 | -7.1 | 0.07 | 93/121 |
| *vha-15* RNAi (adult-only) | 45 | MAH677 | *sid-1; sqIs71[rgef-1p::sid-1+rgef-1p::gfp]* | *vha-15* | 15.7 | 16.8 | -6.6 | 0.01 | 55/118 |
| | 49 | MAH677 | *sid-1; sqIs71[rgef-1p::sid-1+rgef-1p::gfp]* | *vha-15* | 20.3 | 20 | -2.9 | 0.2 | 62/103 |
| *vha-15* RNAi (whole-life) | 50 | MAH677 | *sid-1; sqIs71[rgef-1p::sid-1+rgef-1p::gfp]* | *vha-15* | 18.2 | 18.4 | -1.1 | 0.4 | 99/121 |
| *vha-16* RNAi (adult-only) | 45 | MAH677 | *sid-1; sqIs71[rgef-1p::sid-1+rgef-1p::gfp]* | *vha-16* | 14.5 | 16.8 | -13.7 | 0.0002 | 61/120 |
| | 49 | MAH677 | *sid-1; sqIs71[rgef-1p::sid-1+rgef-1p::gfp]* | *vha-16* | 19.1 | 20 | -8.6 | 0.06 | 61/102 |
| *vha-16* RNAi (whole-life) | 50 | MAH677 | *sid-1; sqIs71[rgef-1p::sid-1+rgef-1p::gfp]* | *vha-16* | 14.5 | 18.4 | -11.2 | <0.0001 | 63/120 |
| *daf-2* RNAi (adult-only) | 51 | MAH676 | *sid-1; sqIs69[rgef-1p::sid-1+rgef-1p::gfp]* | *daf-2* | 27.3 | 17.9 | 48.0 | <0.0001 | 107/120 |
| | 52 | MAH676 | *sid-1; sqIs69[rgef-1p::sid-1+rgef-1p::gfp]* | *daf-2* | 30.1 | 18.1 | 66.2 | <0.0001 | 88/121 |
| | 53 | MAH676 | *sid-1; sqIs69[rgef-1p::sid-1+rgef-1p::gfp]* | *daf-2* | 29.1 | 17.8 | 63.0 | <0.0001 | 63/89 |
| | 5 | MAH677 | *sid-1; sqIs71[rgef-1p::sid-1+rgef-1p::gfp]* | *daf-2* | 39.5 | 18.6 | 112.0 | <0.0001 | 79/153 |
| | 12 | MAH677 | *sid-1; sqIs71[rgef-1p::sid-1+rgef-1p::gfp]* | *daf-2* | 29.1 | 18.3 | 59.1 | <0.0001 | 84/121 |
| | 14 | MAH677 | *sid-1; sqIs71[rgef-1p::sid-1+rgef-1p::gfp]* | *daf-2* | 27.9 | 18.7 | 49.2 | <0.0001 | 58/105 |
| | 5 | MAH757 | *sid-1; sqEx101[rgef-1p::sid-1 + rgef-1p::gfp]* | *daf-2* | 28.7 | 17.3 | 65.8 | <0.0001 | 80/122 |
| | 45 | MAH677 | *sid-1; sqIs71[rgef-1p::sid-1+rgef-1p::gfp]* | *daf-2* | 29.3 | 16.8 | 74.4 | <0.0001 | 56/120 |
| *daf-2* RNAi (whole-life) | 22 | MAH677 | *sid-1; sqIs71[rgef-1p::sid-1+rgef-1p::gfp]* | *daf-2* | 31.5 | 19.9 | 58.2 | <0.0001 | 85/121 |
| | 54 | MAH798 | *sid-1; Ex rgef-1p::sid-1+unc-122p::rfp* | *daf-2* | 32.6 | 17.9 | 82.1 | <0.0001 | 66/120 |
| | 55 | MAH798 | *sid-1; Ex rgef-1p::sid-1+unc-122p::rfp* | *daf-2* | 30.7 | 18.3 | 67.8 | <0.0001 | 69/105 |
| *daf-2* RNAi (L1) | 39 | MAH676 | *sid-1; sqIs69[rgef-1p::sid-1+rgef-1p::gfp]* | *daf-2* | 20.1 | 18.5 | 9.0 | 0.04 | 71/83 |
| | 40 | MAH677 | *sid-1; sqIs71[rgef-1p::sid-1+rgef-1p::gfp]* | *daf-2* | 27.1 | 18.1 | 50.0 | <0.0001 | 97/120 |
| | 41 | MAH676 | *sid-1; sqIs69[rgef-1p::sid-1+rgef-1p::gfp]* | *daf-2(L4)* | 24.4 | 16.8 | 45.0 | <0.0001 | 40/60 |

**Supplementary Table 2:** Lifespan analysis of neuronal-only RNAi strains (Strain name and genotype indicated). RNAi treatment against indicated autophagy genes from indicated time of onset. Exp: experiment number (if multiple RNAi clones were tested in the same experiment, the experiment has the same EXP number also in **Tables S4 and S5**. MLS: mean lifespan; % MLS: percentage change in lifespan compared with control; *P*-values calculated by log-rank test. N: observed deaths/total number of animals. * data depicted in **Figure 1e.**

**Table S3: Summary of lifespan experiments in transgenic strains after neuronal knockdown of *lgg-1* and *atg-7***

| Strain | Exp # | Strain Name | Strain Genotype | RNAi treat-ment | RNAi MLS (Days) | Control MLS (Days) | % MLS change | P value | N animals/ total |
|---|---|---|---|---|---|---|---|---|---|
| *rgef-1p::Q40::yfp* | 1* | MAH828 | *sid-1(qt9) V; rgef-1p::sid-1 + unc-122p::rfp; rgef-1p::Q40::yfp* | *lgg-1* | 14.8 | 11.2 | 32 | <0.0001 | 89/113 |
| | 2 | MAH828 | *sid-1(qt9) V; rgef-1p::sid-1 + unc-122p::rfp; rgef-1p::Q40::yfp* | *lgg-1* | 12.8 | 10.4 | 23 | 0.003 | 99/105 |
| | 3 | MAH828 | *sid-1(qt9) V; rgef-1p::sid-1 + unc-122p::rfp; rgef-1p::Q40::yfp* | *lgg-1* | 19.8 | 18.1 | 9 | 0.009 | 71/96 |
| | 1* | MAH828 | *sid-1(qt9) V; rgef-1p::sid-1 + unc-122p::rfp; rgef-1p::Q40::yfp* | *atg-7* | 13.7 | 11.2 | 22 | <0.0001 | 80/102 |
| | 2 | MAH828 | *sid-1(qt9) V; rgef-1p::sid-1 + unc-122p::rfp; rgef-1p::Q40::yfp* | *atg-7* | 14.6 | 10.4 | 41 | <0.0001 | 95/105 |
| | 3 | MAH828 | *sid-1(qt9) V; rgef-1p::sid-1 + unc-122p::rfp; rgef-1p::Q40::yfp* | *atg-7* | 17.4 | 18.1 | -4 | 0.5 | 79/96 |
| *mec-4p::mCherry* | 4 | ZB5184 | *sid-1(qt9) V; mec-4p::mCherry; rgef-1p::gfp + rgef-1p::sid 1 + pBS* | *lgg-1* | 37.9 | 29.0 | 30 | <0.0001 | 65/90 |
| | 5 | ZB5184 | *sid-1(qt9) V; mec-4p::mCherry; rgef-1p::gfp + rgef-1p::sid 1 + pBS* | *lgg-1* | 30.0 | 24.3 | 23 | <0.0001 | 74/90 |
| | 6 | ZB5184 | *sid-1(qt9) V; mec-4p::mCherry; rgef-1p::gfp + rgef-1p::sid 1 + pBS* | *lgg-1* | 27.7 | 23.1 | 30 | <0.0001 | 63/96 |
| | 4 | ZB5184 | *sid-1(qt9) V; mec-4p::mCherry; rgef-1p::gfp + rgef-1p::sid 1 + pBS* | *atg-7* | 31.5 | 29.0 | 8 | 0.003 | 52/90 |
| | 5 | ZB5184 | *sid-1(qt9) V; mec-4p::mCherry; rgef-1p::gfp + rgef-1p::sid 1 + pBS* | *atg-7* | 29.1 | 24.3 | 19 | 0.0001 | 78/90 |
| | 6 | ZB5184 | *sid-1(qt9) V; mec-4p::mCherry; rgef-1p::gfp + rgef-1p::sid 1 + pBS* | *atg-7* | 25.1 | 23.1 | 8 | 0.06 | 63/96 |

**Supplementary Table 3**: Lifespan analysis of neuronal-only RNAi strains expressing neuronal Q40 or mCherry in mechano-sensory neurons (Strain name and genotype indicated). RNAi treatment against indicated autophagy genes from hatching on. Exp: experiment number (if multiple RNAi clones were tested in the same experiment, the experiment has the same EXP number in this Table). MLS: mean lifespan; % MLS: percentage change in lifespan compared with control; P-values calculated by log-rank test. N: observed deaths/total number of animals. * data depicted in **Figure S2c.**

**Table S4: Summary of lifespan experiments in autophagy mutants *atg-4.1* and *atg-16.2***

| Strain / Comment | Exp # | Strain Name | Strain Genotype | RNAi treat-ment | RNAi MLS (Days) | Control MLS (Days) | % MLS change | P value | N animals/ total |
|---|---|---|---|---|---|---|---|---|---|
| *atg-4.1(bp501)* | 30 | MAH977 | *atg-4.1; sid-1; rgef-1p::sid-1 + unc-122p::rfp* | *lgg-1* | 16.1 | 14.3 | 12.6 | 0.006 | 71/119 |
| | 30 | MAH977 | *atg-4.1; sid-1; rgef-1p::sid-1 + unc-122p::rfp* | *atg-7* | 15.8 | 14.3 | 10.5 | 0.01 | 62/115 |
| | 31 | MAH977 | *atg-4.1; sid-1; rgef-1p::sid-1 + unc-122p::rfp* | *lgg-1* | 18.7 | 16.2 | 15.4 | <0.0001 | 93/147 |
| | 31 | MAH977 | *atg-4.1; sid-1; rgef-1p::sid-1 + unc-122p::rfp* | *atg-7* | 18.7 | 16.2 | 15.4 | <0.0001 | 110/160 |
| | 32* | MAH977 | *atg-4.1; sid-1; rgef-1p::sid-1 + unc-122p::rfp* | *lgg-1* | 19.7 | 15.6 | 26.3 | <0.0001 | 102/140 |
| | 32* | MAH977 | *atg-4.1; sid-1; rgef-1p::sid-1 + unc-122p::rfp* | *atg-7* | 18.2 | 15.6 | 16.7 | <0.0001 | 99/140 |
| *atg-16.2(ok3224)* | 26 | MAH899 | *atg-16.2; sid-1; rgef-1p::sid-1 + rgef-1p::gfp* | *lgg-1* | 18.9 | 18.8 | 0.5 | 0.9 | 108/120 |
| | 26 | MAH980 | *atg-16.2; sid-1; rgef-1p::sid-1 + unc-122p::rfp* | *atg-7* | 18.6 | 18.8 | -1.1 | 1 | 100/121 |
| | 27 | MAH899 | *atg-16.2; sid-1; rgef-1p::sid-1 + rgef-1p::gfp* | *lgg-1* | 19.3 | 18.7 | 3.2 | 0.2 | 82/120 |
| | 27 | MAH980 | *atg-16.2; sid-1; rgef-1p::sid-1 + unc-122p::rfp* | *atg-7* | 18.1 | 18.7 | -3.3 | 0.7 | 100/120 |
| | 30* | MAH980 | *atg-16.2; sid-1; rgef-1p::sid-1 + unc-122p::rfp* | *lgg-1* | 15.6 | 15.2 | 2.6 | 0.6 | 85/117 |
| | 30* | MAH980 | *atg-16.2; sid-1; rgef-1p::sid-1 + unc-122p::rfp* | *atg-7* | 14.9 | 15.2 | 2.1 | 0.5 | 84/120 |
| | 54 | MAH980 | *atg-16.2; sid-1; rgef-1p::sid-1 + unc-122p::rfp* | *daf-2* | 29.7 | 15.2 | 95.4 | <0.0001 | 59/115 |
| | 45 | MAH980 | *atg-16.2; sid-1; rgef-1p::sid-1 + unc-122p::rfp* | *daf-2* | 23.5 | 15.9 | 95.4 | <0.0001 | 59/115 |
| | 55 | MAH980 | *atg-16.2; sid-1; rgef-1p::sid-1 + unc-122p::rfp* | *daf-2* | 24.1 | 16.8 | 50.3 | <0.0001 | 75/120 |

**Supplementary Table 4**: Lifespan analysis of *atg-4.1(bp501)* and *atg-16.2(ok3224)* mutants capable of neuronal-only RNAi (Strain name and genotype indicated). RNAi treatment against indicated autophagy genes from hatching on. Exp: experiment number (Control in WT animals listed with same Exp# in **Table S2**) MLS: mean lifespan; % MLS: percentage change in lifespan compared with control;  P-values calculated by log-rank test. N: observed deaths/total number of animals. * data depicted in **Figure 4e**.

**Table S5: Summary of lifespan experiments with neuronal *atg-16.2* reconstitution**

| Strain / Comment | Exp # | Strain Name | Strain Genotype | RNAi treat-ment | RNAi MLS (Days) | Control MLS (Days) | % MLS change | P value | N animals/ total |
|---|---|---|---|---|---|---|---|---|---|
| *atg-16.2; rgef-1p::atg-16.2* | 33 | MAH1036 | *atg-16.2; sid-1; rgef-1p::sid-1 + unc-122p::rfp; rgef-1p::atg-16.2 + rol-6* | *lgg-1* | 19.1 | 13.7 | 39.4 | <0.0001 | 59/100 |
| | | | | *atg-7* | 16.3 | 13.7 | 18.9 | 0.003 | 62/113 |
| | 34* | MAH1036 | *atg-16.2; sid-1; rgef-1p::sid-1 + unc-122p::rfp; rgef-1p::atg-16.2 + rol-6* | *lgg-1* | 18.3 | 14.7 | 24.5 | <0.0001 | 70/109 |
| | | | | *atg-7* | 20.6 | 14.7 | 40.1 | <0.0001 | 65/120 |
| | 35 | MAH1036 | *atg-16.2; sid-1; rgef-1p::sid-1 + unc-122p::rfp; rgef-1p::atg-16.2 + rol-6* | *lgg-1* | 15.5 | 14.1 | 9.9 | 0.08 | 60/102 |
| | | | | *atg-7* | 15.8 | 14.1 | 12.1 | 0.03 | 65/100 |
| *atg-16.2; rgef-1p::atg-16.2ΔWD40* | 32* | MAH1025 | *atg-16.2; sid-1; rgef-1p::sid-1 + unc-122p::rfp; rgef-1p::atg-16.2ΔWD40 + rol-6* | *lgg-1* | 15.6 | 15.1 | 3.3 | 0.5 | 60/89 |
| | | | | *atg-7* | 14.5 | 15.1 | -4.0 | 0.5 | 68/97 |
| | 33 | MAH1025 | *atg-16.2; sid-1; rgef-1p::sid-1 +unc-122p::rfp; rgef-1p::atg-16.2ΔWD40 + rol-6* | *lgg-1* | 13.7 | 14.0 | -2.2 | 0.2 | 80/122 |
| | | | | *atg-7* | 13.8 | 14.0 | -1.5 | 0.04 | 75/120 |
| | 33 | MAH1024 | *atg-16.2; sid-1; rgef-1p::sid-1 +unc-122p::rfp; rgef-1p::atg-16.2ΔWD40 + rol-6* | *lgg-1* | 14.1 | 14.5 | -2.8 | 0.5 | 61/98 |
| | | | | *atg-7* | 15.1 | 14.5 | 4.1 | 0.8 | 71/120 |
| *atg-16.2; rgef-1p::atg-16.2 (F394A)* | 33 | MAH1038 | *atg-16.2; sid-1; rgef-1p::sid-1 + unc-122p::rfp; rgef-1p::atg-16.2(F394A) + rol-6* | *lgg-1* | 14.1 | 14.5 | -2.8 | 0.4 | 33/60 |
| | | | | *atg-7* | 15.1 | 14.5 | 4.1 | 0.9 | 27/46 |
| | 34* | MAH1038 | *atg-16.2; sid-1; rgef-1p::sid-1 + unc-122p::rfp; rgef-1p::atg-16.2(F394A) + rol-6* | *lgg-1* | 16.9 | 16.5 | 2.4 | 0.5 | 67/120 |
| | | | | *atg-7* | 15.8 | 16.5 | -4.3 | 0.4 | 61/120 |
| | 35 | MAH1038 | *atg-16.2; sid-1; rgef-1p::sid-1 + unc-122p::rfp; rgef-1p::atg-16.2(F394A) + rol-6* | *lgg-1* | 14.4 | 15.2 | -5.3 | 0.4 | 33/61 |
| | | | | *atg-7* | 13.7 | 15.2 | -5.9 | 0.06 | 63/105 |

**Supplementary Table 5:** Lifespan analysis of *atg-16.2(ok3224)* mutants capable of neuronal-only RNAi with neuronal reconstitution of ATG-16.2 variants (Strain name and genotype indicated). RNAi treatment against indicated autophagy genes from hatching on. Exp: experiment number (Control in WT animals listed with same Exp# in **Table S2**) MLS: mean lifespan; % MLS: percentage change in lifespan compared with control; P-values calculated by log-rank test. N: observed deaths/total number of animals. * data depicted in **Figure 6g-i**.

**Table S6: Strains used in this study**

| Comment | Strain | Genotype |
|---|---|---|
| published | N2 - Hansen | *Wild-type (WT)* |
| | AM101 | *rmIs110[rgef-1p::Q40::YFP]* |
| | CB1387 | *daf-10(e1387) IV* |
| | MAH242 | *sqIs24[rgef-1p::gfp::lgg-1 + unc-122p::rfp]* |
| | MAH346 | *sid-1(qt9) V* |
| | MAH349 | *sqIs35[pMH951/sqst-1p::sqst-1::gfp+pMH876/unc-122p::rfp]* |
| | PR811 | *osm-6(p811) V* |
| | XE1375 | *wpIs36[unc-47p::mCherry] I; wpSi1[unc-47p::rde-1::SL2::sid-1 + unc-119(+)] II; eri-1(mg366) IV; rde-1(ne219) V; lin-15B(n744) X* |
| | ZB4065 | *bzIs166[mec-4p::mCherry]* |
| newly generated | MAH629 | *sqEx100[rgef-1p::gfp + pBS]* |
| | MAH630 | *sqEx101[rgef-1p::gfp + rgef-1p::sid-1 + pBS]* |
| | MAH656 | *sqIs69[rgef-1p::gfp + rgef-1p::sid-1 + pBS]* |
| | MAH657 | *sqIs71[rgef-1p::gfp + rgef-1p::sid-1 + pBS]* |
| | MAH676 | *sid-1(qt9) V; sqIs69[rgef-1p::gfp + rgef-1p::sid-1 + pBS]* |
| | MAH677 | *sid-1; sqIs71[rgef-1p::sid-1+ rgef-1p::gfp + pBS]* |
| | MAH732 | *atg-4.1(bp501) I; sqIs24[rgef-1p::gfp::lgg-1 + unc-122p::rfp]* |
| | MAH752 | *sid-1(qt9) V; rmIs110 [rgef-1p::Q40::yfp]* |
| | MAH757 | *sid-1(qt9) V; sqEx101[rgef-1p::gfp + rgef-1p::sid-1 + pBS]* |
| | MAH798 | *sid-1(qt9) V; sqEx131[rgef-1p::sid-1 + unc-122p::rfp]* |
| | MAH810 | *sid-1(qt9) V; sqIs24[rgef-1p::gfp::lgg-1 + unc-122p::rfp]* |
| | MAH828 | *sid-1(qt9) V; sqEx131[rgef-1p::sid-1 + unc-122p::rfp]; rmIs110 [rgef-1p::Q40::yfp]* |
| | MAH848 | *sid-1(qt9) V; sqEx148[rgef-1p::sid-1 + rol-6]* |
| | MAH899 | *atg-16.2(ok3224) II; sid-1(qt9) V; sqIs71[rgef-1p::gfp + rgef-1p::sid-1 +pBS]* |
| | MAH917 | *sqIs90[rgef-1p::gfp::lgg-1(G116A) + unc-122p::rfp]* |
| | MAH923 | *sid-1(qt9) V; sqIs24[rgef-1p::gfp::lgg-1 + unc-122p::rfp]; sqEx148[rgef-1p::sid-1 +rol-6]* |
| | MAH973 | *atg-16.2(ok3224) II* |
| | MAH977 | *atg-4.1(bp501) I; sid-1(qt9) V; sqEx131[rgef-1p::sid-1 + unc-122p::rfp]* |
| | MAH979 | *atg-16.2(ok3224) II; sid-1(qt9) V* |

| | | |
|---|---|---|
| | MAH980 | *atg-16.2(ok3224) II; sid-1(qt9) V; sqEx131[rgef-1p::sid-1 + unc-122p::rfp]* |
| | MAH981 | *atg-4.1(bp501) I; sid-1(qt9) V; rmIs110 [rgef-1p::Q40::yfp]; sqEx131[rgef-1p::sid-1 + unc-122p::rfp]* |
| | MAH1024 | *atg-16.2(ok3224) II; sid-1(qt9) V; sqEx131[rgef-1p::sid-1 + unc-122p::rfp]; sqEx171[rgef-1p::atg-16.2ΔWD40 + rol-6]* |
| | MAH1025 | *atg-16.2(ok3224) II; sid-1(qt9) V; sqEx131[rgef-1p::sid-1 + unc-122p::rfp]; sqEx172[rgef-1p::atg-16.2ΔWD40 + rol-6]* |
| | MAH1027 | *atg-16.2(ok3224) II; sid-1(qt9) V; sqEx131[rgef-1p::sid-1 + unc-122p::rfp]; rmIs110 [rgef-1p::Q40::yfp]* |
| | MAH1036 | *atg-16.2(ok3224) II; sid-1(qt9) V; sqEx131[rgef-1p::sid-1 + unc-122p::rfp]; sqEx173[rgef-1p::atg-16.2 + rol-6]* |
| | MAH1038 | *atg-16.2(ok3224) II; sid-1(qt9) V; sqEx131[rgef-1p::sid-1 + unc-122p::rfp]; sqEx175[rgef-1p::atg-16.2(F394A) + rol-6]* |
| | MAH1051 | *atg-16.2(ok3224) II; sqIs92[rgef-1p::gfp::lgg-1(G116A) + unc-122p::rfp]* |
| | MAH1052 | *atg-4.1(bp501) I; sqIs92[rgef-1p::gfp::lgg-1(G116A) + unc-122p::rfp]* |
| | MAH1053 | *atg-16.2(ok3224) II; sid-1(qt9) V; rmIs110 [rgef-1p::Q40::yfp]; sqEx131[rgef-1p::sid-1 + unc-122p::rfp]; sqEx173 [rgef-1p::atg-16.2 + rol-6]* |
| | MAH1054 | *atg-16.2(ok3224) II; sid-1(qt9) V; rmIs110 [rgef-1p::Q40::yfp]; sqEx131[rgef-1p::sid-1 + unc-122p::rfp]; sqEx172[rgef-1p::atg-16.2ΔWD40 + rol-6]* |
| | MAH1055 | *atg-16.2(ok3224) II; sid-1(qt9) V; rmIs110 [rgef-1p::Q40::yfp]; sqEx131[rgef-1p::sid-1 + unc-122p::rfp]; sqEx175[rgef-1p::atg-16.2(F394A) + rol-6]* |
| | MAH1126 | *atg-4.1(bp501) I* |
| | MAH1186 | *sid-1(qt9) V; sqIs35[sqst-1p::sqst-1::gfp + unc-122p::rfp]* |
| | MAH1194 | *sid-1(qt9) V; sqIs35[sqst-1p::sqst-1::gfp + unc-122p::rfp]; sqEx148[rgef-1p::sid-1 + rol-6]* |
| | MAH1195 | *atg-16.2(ok3224) II; sqIs24[rgef-1p::gfp::lgg-1 + unc-122p::rfp]* |
| | MAH1198 | *atg-16.2(ok3224) II; sqIs35[sqst-1p::sqst-1::gfp + unc-122p::rfp]* |
| | MAH1209 | *sid-1(qt9) V; sqIs90[gef-1p::gfp::lgg-1(G116A) +unc-122p::rfp]* |
| | MAH1212 | *atg-4.1(bp501) I; sqIs35[sqst-1p::sqst-1::gfp + unc-122p::rfp]* |
| | MAH1215 | *sid-1(qt9) V; sqIs90[rgef-1p::gfp::lgg-1(G116A) + unc-122p::rfp]; sqEx148[rgef-1p::sid-1 + rol-6]* |
| | MAH1230 | *sid-1(qt9) V; bzIs166[mec-4p::mCherry]* |
| | KUM7 | *atg-16.2(ok3224) II; sid-1(qt9) V; bzIs166[mec-4p::mCherry]; sqIs71[rgef-1p::gfp + rgef-1p::sid 1 + pBS]* |
| | KUM16 | *atg-4.1(bp501) I; bzIs166[mec-4p::mCherry]* |
| | KUM19 | *bzIs166[mec-4p::mCherry]; bzEx305[rgef-1p::atg-16.2 + pCFJ1662(HygR)]* |
| | KUM20 | *bzIs166[mec-4p::mCherry]; bzEx302[rgef-1p::atg-16.2ΔWD40 + pCFJ1662(HygR)]* |
| | KUM28 | *atg-16.2(ok3224) II; bzIs166[mec-4p::mCherry]; bzEx305[rgef-1p::atg-16.2 + pCFJ1662(HygR)]* |
| | KUM40 | *atg-16.2(ok3224) II; bzIs166[mec-4p::mCherry]; bzEx302[rgef-1p::atg-16.2ΔWD40 + pCFJ1662(HygR)]* |
| | KUM43 | *atg-16.2(ok3224) II; bzIs166[mec-4p::mCherry]; bzEx305[rgef-1p::atg-16.2 + pCFJ1662(HygR)], sqIs71[rgef-1p::gfp + rgef-1p::sid-1 + pBS]* |
|  | KUM46 | *atg-16.2(ok3224) II; sid-1(qt9) V; bzIs166[mec-4p::mCherry]; bzEx300[rgef-1p::atg-16.2ΔWD40 + pBG33H6(HygR)]* |

| | | |
|---|---|---|
| | KUM48 | *atg-16.2(ok3224) II; bzIs166[mec-4p::mCherry]; bzEx301[rgef-1p::atg-16.2(F394A) + pCFJ1662(HygR)]* |
| | KUM49 | *atg-16.2(ok3224) II; sid-1(qt9) V; bzIs166[mec-4p::mCherry]; sqIs71[rgef-1p::gfp + rgef-1p::sid 1 + pBS]; bzEx300[rgef-1p::atg-16.2∆WD40 + pBG33H6(HygR)]* |
| | KUM50 | *atg-16.2(ok3224) II; sid-1(qt9) V; bzIs166[mec-4p::mCherry]; sqIs71[rgef-1p::gfp + rgef-1p::sid 1 + pBS]; bzEx301[rgef-1p::atg-16.2(F394A) + pCFJ1662(HygR)]* |
| | KUM51 | *atg-16.2(ok3224) II; sqIs24[rgef-1p::gfp::lgg-1 + unc-122p::rfp]; bzEx301[rgef-1p::atg-16.2(F394A) + pCFJ1662(HygR)]* |
| | KUM55 | *atg-16.2(ok3224) II; sqIs24[rgef-1p::gfp::lgg-1 + unc-122p::rfp]; bzEx302[rgef-1p::atg-16.2∆WD40 + pCFJ1662(HygR)]* |
| | KUM57 | *atg-16.2(ok3224) II; sqIs24[rgef-1p::gfp::lgg-1 + unc-122p::rfp]; bzEx305[rgef-1p::atg-16.2 + pCFJ1662(HygR)]* |
| | KUM59 | *atg-16.2(ok3224) II; sid-1(qt9) V; bzIs166[mec-4p::mCherry]; sqIs71[rgef-1p::gfp + rgef-1p::sid 1 + pBS]; bzEx305[rgef-1p::atg-16.2 + pCFJ1662(HygR)]* |
| | ZB5167 | *atg-16.2(ok3224) II; bzIs166[mec-4p::mCherry]* |
| | ZB5174 | *bzEx302[rgef-1p::atg-16.2∆WD40 + pCFJ1662(HygR)]* |
| | ZB5184 | *sid-1(qt9) V; bzIs166[mec-4p::mCherry]; sqIs71[rgef-1p::gfp + rgef-1p::sid 1 + pBS]* |
| | ZB5329 | *bzIs166[mec-4p::mCherry]; bzEx301[rgef-1p::atg-16.2(F394A) + pCFJ1662(HygR)]* |
| | ZB5330 | *bzEx300[rgef-1p::atg-16.2∆WD40 + pBG33H6(HygR)]* |
| | ZB5333 | *bzEx305[rgef-1p::atg-16.2 + pCFJ1662(HygR)]* |

**Supplementary Table 6:** Strains used in this study. Strain name and genotype indicated.

**Table S7: Plasmids used in this study**

| Comment | Plasmid Name | Plasmid description |
|---|---|---|
| published | pBluescript | empty vector - used as "stuffer" DNA in injections to bring total DNA concentration up to 100 ng/μl |
| | pCFJ1662 | Hygromycin resistance |
| | pMH876 | *unc-122p::rfp* |
| | pRF4 | *rol-6(su1006)* |
| newly generated | pBG33H6 | Hygromycin resistance, pCFJ1662 with Gateway cloning sites inserted |
| | pMH1141 | *rgef-1p::sid-1* - Gateway destination vector pDEST_R3R4 with *rgef-1* pan-neuronal promoter, *sid-1* ORF, and *unc-54* 3'-UTR |
| | pMH1201 | *rgef-1p::gfp* - Gateway destination vector pDEST_R3R4 with *rgef-1 pan-neuronal* promoter, *gfp* ORF, and *unc-54* 3'-UTR |
| | pMH1355 | L4440(-114 bp); RNAi empty vector control - L4440 digested with EcoRV and re-ligated |
| | pMH1387 | *rgef-1p::atg-16* |
| | pMH1388 | *rgef-1p::atg-16(F394A)* |
| | pMH1389 | *rgef-1p::atg-16ΔWD40* |
| RNAi clones | L4440 | Dr. Andrew Fire |
| | L4440 (EcoRV-digested) | this study |
| | *lgg-1* | JA |
| | *atg-7* | JA |
| | *bec-1* | JA |
| | *atg-9* | JA |
| | *atg-13* | JA |
| | *unc-51* | MV |
| | *atg-4.1* | MV |
| | *atg-16.2* | MV |
| | *epg-5* | JA |
| | *cup-5* | JA |
| | *vha-13* | JA |
| | *vha-15* | JA |
| | *vha-16* | MV |
| | *daf-2* | Dr. Andrew Dillin |
| | *gfp* | Dr. Andrew Dillin |

| | | |
|---|---|---|
| | *snb-1* | MV |
| | *snb-1* | JA |
| | *unc-13* | JA |
| | *elt-2* | JA |
| | *pept-1* | JA |
| | *bli-3* | JA |
| | *die-1* | JA |
| | *unc-112* | JA |
| | *unc-22* | JA |
| | *tsp-15* | Dr. Hiroki Mirobe |
| | *rpl-2* | JA |

**Supplementary Table 7:** Plasmids used in this study. Published and newly generated plasmids are indicated and fully described in strain construction. RNAi clones indicated with origin. JA: Julie Ahringer RNAi library, MV: Marc Vidal RNAi library. Some RNAi clones were gifts.

**Table S8: Primers used in this study**

| Comment | Name | | Sequence 5'-3' | Used for |
|---|---|---|---|---|
| plasmid generation | *atg-16.2ΔWD40* | FWD | CGGCTAGCATGACCGACAATCGGACTTC | To generate pMH1389 by Gibson assembly of *atg-16.2delC* fragment containing Stop codon into plasmid containing *rgef-1p::gfp* |
| | *atg-16.2ΔWD40*-STOP | REV | GCGTCGACTCAAGTTTCGGGAGATGTACCAA | |
| | *atg-16.2* | FWD | CAGGAGGACCCTTGGCTAGCATGACCGACAATCGGACTTC | To generate pMH1387 by Gibson assembly of full-length *atg-16.2* cDNA into plasmid containing *rgef-1p::gfp* |
| | *atg-16.2* | REV | AATACCATGGTACCGTCGACTCATCTCCAGAGTGTACAAG | |
| | *atg-16.2(F394A)* | FWD | TCTTCCCATGCTGATAAAAAGG | To generate pMH1388 by site-directed mutagenesis of pMH1387 |
| | *atg-16.2(F394A)* | REV | AATGAAGCTGGATTGAGAAACA | |
| qRT-PCR | *atg-16.2(before ok3224)* | FWD | GGCTGACAGTGAATCTCGTTATTC | *atg-16.2* expression levels for all constructs |
| | *atg-16.2 (before ok3224)* | REV | CTTTCAATTCGGCCAACTCGTT | *atg-16.2* expression levels for all constructs |
| | *atg-16.2 (in WD40)* | FWD | GACTATTGAGTACGTTTTCTGGCC | *atg-16.2* expression levels for full-length and F394A constructs |
| | *atg-16.2(in WD40)* | REV | CCCAATTCTTGATTGTACGATCCGC | *atg-16.2* expression levels for full-length and F394A constructs |
| | *cyn-1* | FWD | GTGTCACCATGGAGTTGTTC | Housekeeping gene |
| | *cyn-1* | REV | TCCGTAGATTGATTCACCAC | Housekeeping gene |
| | *pmp-3* | FWD | GTTCCCGTGTTCATCACTCAT | Housekeeping gene |
| | *pmp-3* | REV | ACACCGTCGAGAAGCTGTAGA | Housekeeping gene |
| | *nhr-23* | FWD | CAGAAACACTGAAGAACGCG | Housekeeping gene |
| | *nhr-23* | REV | CGATCTGCAGTGAATAGCTC | Housekeeping gene |

**Supplementary Table 8:** Primers used for plasmid generation and qRT-PCR. All Sequences indicated in 5'-3'.
